# Septum site placement in *Mycobacteria* - Identification and Characterization of mycobacterial homologues of *Escherichia coli* MinD

**DOI:** 10.1101/2023.03.20.533423

**Authors:** Vimal Kishore, Sujata S. Gaiwala Sharma, Tirumalai R. Raghunand

## Abstract

A major virulence trait of *Mycobacterium tuberculosis* (*M. tb*) is its ability to enter a dormant state within its human host. Since cell division is intimately linked to metabolic shut down, understanding the mechanism of septum formation and its integration with other events in the division pathway is likely to offer clues to the molecular basis of dormancy. The *M. tb* genome lacks obvious homologues of several conserved cell division proteins, and this study aimed at identifying and functionally characterising mycobacterial homologues of the *E.coli* septum site specification protein MinD (*Ec* MinD). Sequence homology based analyses suggested that the genomes of both *M.tb* and the saprophyte *Mycobacterium smegmatis* (*M. smegmatis*) encode two putative *Ec* MinD homologues *-* Rv1708/MSMEG_3743 and Rv3660c/MSMEG_6171. Both *Rv1708* and *MSMEG_3743* were observed to fully complement the mini-cell phenotype of the *E.coli* Δ*minDE* mutant *HL1,* but the other homologues only partially complemented the mutant phenotype. Over-expression of *MSMEG_3743* but not *MSMEG_6171* in *M. smegmatis* led to cell elongation and a drastic decrease in CFU counts, indicating the essentiality of *MSMEG_3743* in cell-division. Sequence analysis of MSMEG_3743 showed a conserved Walker A motif, the functional role of which was confirmed by a radiolabelled ATPase activity assay. Rv1708 was observed to interact with the chromosome associated proteins ScpA and ParB, pointing to a link between its septum formation role and chromosome segregation. Comparative structural analyses showed Rv1708 to be closer in similarity to *Ec* MinD than Rv3660c. In summary we have demonstrated that Rv1708 and MSMEG_3743 are true mycobacterial homologues of *Ec* MinD, adding a critical missing piece to the mycobacterial cell division puzzle.

## INTRODUCTION

Cell division is crucial to all living organisms and all cells have developed mechanisms for the equal distribution of genetic and cytoplasmic material during the process of division. In bacteria, cell division begins from marking the division site, usually at the mid-cell. This is largely determined by two negative regulatory mechanisms - the Min System and Nucleoid Occlusion system, which ensure inhibition of FtsZ polymerization at the poles, and decapitating the bacterial chromosome respectively. The Min system in *E. coli* consists of three genes - *minC, minD* and *minE* encoded in the *minB* operon, first described due to its mini-cell phenotype [1, 2], which results from polar localization of the FtsZ ring. MinD directly binds to the H10 helix of FtsZ with a higher affinity than MinC-FtsZ, and triggers MinC activity to inhibit FtsZ functions [3]. The system maintains cell size by recycling FtsZ and inhibits peptidoglycan turnover especially at the cell poles. This system is involved in swarming of *Proteus mirabilis* [4], and is also known to affect the localization of metabolic enzymes to the *E.coli* inner membrane [5]. In addition, A role for MinD has been described in the virulence in *Aeromonas hydrophila* [6], *Neisseria gonorrhoeae* [7], *Helicobacter pylori* [8], and *Listeria monocytogenes* [9]. MinD is one of the major members of the MinD/ParA/SojA WACA (Walker A Cytoskeletal ATPase) family. It is an amphitropic protein and localizes to the cytoplasm or membrane depending on substrate binding [10]. The protein has a conserved Membrane Targeting Sequence by which it binds to the membrane in an ATP dependent manner and this sequence is highly conserved from bacteria to chloroplasts [11]. In addition, MinD contains the ATPase associated Deviant Walker A motif and displays ATPase activity [12, 13].

Several components of the cell-division machinery identified in *E. coli* and *B. subtilis* are conserved in mycobacteria [14], however, a majority of them are still un-annotated and uncharacterized. In mycobacteria, the placement of the FtsZ ring at the mid-cell is not as stringent as in *E. coli, B. subtilis* or *C. glutamicum*, with the characteristic feature of mycobacterial cell-division being the formation of the asymmetric V-shape during cytokinesis [14, 15]. Two models explain the asymmetric nature of mycobacterial division - both of which invoke the faster growth of daughter cells containing the old pole, and differ only in their description of whether asymmetry arises due to faster growth of these cells either before or after cytokinesis [16–18].

Since no Min like system has been reported in mycobacteria so far, we set out to identify such MinD/ParA/SojA family members, to gain insights into the process of mycobacterial septum site determination, using a target-based genetic complementation approach [19]. Following the identification of MSMEG_3743 and Rv1708 as homologues of *Ec* MinD, we proceeded to carry out a detailed characterization of the *M. smegmatis* MinD homologue, MSMEG_3743 and investigated its role in mycobacterial cell division using genetic, biochemical, cell-biological, and comparative structural analysis approaches. Our findings indicate that in addition to sharing some of the properties described for *E. coli* MinD, MSMEG_3743 possesses some unique features, which may be a reflection of its role in the atypical mechanism of cell division in mycobacteria.

## MATERIALS AND METHODS

### Bacterial culture, Media and Growth Conditions

*E. coli* cultures were grown in LB medium at 37°C with agitation at 200 rpm or on LB agar plates. The *M. smegmatis* strain mc^2^155 was grown in Middlebrook 7H9 broth (Difco) or on Middlebrook 7H10 agar (Difco) supplemented with 10% albumin-dextrose-saline (ADS), 0.2 % glycerol, and 0.05 % Tween-80 at 37°C with agitation at 200 rpm. *M. tb* H37Ra was grown under the same conditions as *M. smegmatis* with the exception of 10% (v/v) Oleic Acid-Albumin-Dextrose-Catalase (OADC) supplementation instead of 10% ADS. Antibiotics were added at the following concentrations: ampicillin, 200 µg/mL, chloramphenicol , 25 µg/mL, tetracycline, 10 µg/mL, kanamycin (50 µg/mL for *E. coli,* 15 µg/mL for *M. smegmatis*), hygromycin (200 µg/mL for *E. coli*, 50 µg/mL for *M. smegmatis*).

### DNA Techniques

Restriction enzymes, Taq polymerase and T4 DNA ligase were purchased from New England Biolabs (NEB). Standard protocols were followed for DNA manipulation, including plasmid DNA preparation, restriction endonuclease digestion, agarose gel electrophoresis, isolation and ligation of DNA fragments, and *E. coli* transformation. DNA fragments used for cloning reactions were purified by using the Nucleospin gel extraction kit (Macherey-Nagel) according to the manufacturer’s specifications. Mycobacterial strains were transformed by electroporation at 1800 V, 1000 µF, and 25 Ω on a Bio-Rad Gene pulser X-cell electroporator.

### Complementation Assays

For complementation assays, ORFs corresponding to *E. coli minD* and its *M. smegmatis* & *M. tb* H37Rv homologues, were amplified from their respective gDNA using the primers listed in Table S1, and cloned into pTrc99A. Recombinant pTrc99A constructs were transformed into the *E. coli* HL1 (Δ*minDE*) mutant strain along with an empty vector as a negative control. Transformed strains were induced with 0.5 mM IPTG at an OD of 0.4-0.6 at 37°C. After 12 h, the frequency of minicells in each sample was determined using DIC microscopy.

### Overexpression Studies

For overexpression assays, ORFs corresponding to the *M. smegmatis* homologues of *Ec minD* (*MSMEG_3743* and *MSMEG_6171*), *Ms parB*, *Ms scpA* and *Ms scpB* were cloned into pTetO using the primers listed in Table S1. The control vector ΔpTetO was constructed by releasing the insert from pSE100-rbs-RFP by restriction digestion followed by end-filling and ligation. The Recombinant pTetO constructs were transformed into *M. smegmatis* mc^2^155 along with the empty vector for use as the negative control. Transformant cultures were induced at an OD of 0.4-0.5 with 50 ng/mL Anhydrotetracycline (ATc) for 16 h at 37°C. The induced cultures were then processed for CFU counts, microscopy and RNA isolation. Data from three independent experiments was used to analyse the overexpression phenotypes depicted.

### Mycobacterial Protein Fragment Complementation (MPFC) assay

To identify interacting partners for the mycobacterial MinD homologues, putative interacting partners for Rv1708 were predicted using the algorithm of Hegde *et al.* [20], and specific candidates were chosen for further analyses. In order to validate these interactions, the bait (*Rv1708, MSMEG_374*3) and prey (*M. tb ParB*/*Ms ParB* &*M. tb ScpA*/*Ms ScpA*) were cloned in the *E. coli*- Mycobacteria shuttle vectors pUAB400 and pUAB300 respectively, using the primers listed in Table S1. Both sets of recombinant plasmids were co-transformed in *M. smegmatis* mc^2^155 along with the appropriate negative controls, and the interaction assay was performed as described in [21].

### Pull-down Assays

To biochemically validate the positive interactions from the MPFC assay, the ORF corresponding to *Rv1708* was cloned with an N-terminal GST tag in pGEX-6P-1, while ORFs corresponding to *M. tb scpA* and *M. tb parB* were cloned into pET22b with C-terminal 6xHis-tags, using the primers in Table S1. Following protein expression (in *E. coli* BL21DE3), purification and quantification, equimolar amounts of both bait and prey proteins were incubated O/N at 4°C in PBS + 0.1% Triton X-100. These were then incubated with equilibrated GSH beads for 4 h at 4°C. Samples were washed (5 x PBS+0.1% Triton X-100 and 5 x PBS), resolved using SDS-PAGE along with the appropriate controls, and subjected to western blotting using anti-6xHis Antibodies.

### Subcellular localization

To determine its subcellular localization, the ORF of *MSMEG_3743* was cloned with a C-terminal 6xHis-tag fusion in the acetamide inducible vector pSCW35 using the primers in Table S1. Cells were harvested 16 h post induction with 0.2% Acetamide at 37°C, and subjected to sonication (30 s on /30 s off, 4 cycles, 5 min). The pellet (Cell debris) was removed from the supernatant by centrifugation (3000 g, 10 min, 4°C). This supernatant was subjected to another step of centrifugation (30,000 g, 30 min, 4°C) and the pellet (Cell wall) was resuspended in 0.5% CHAPS. The supernatant was immediately subjected to another round of centrifugation (100,000 g, 2 h, 4°C) and the pellet (Cell membrane) was resuspended in 0.5% CHAPS. The final supernatant represents the cytoplasmic fraction. All subcellular fractions were quantified using the BCA method and subjected to Western-blotting using an anti-6xHis antibody.

### Western Blotting

To prepare cell lysates, *M. smegmatis* cells were harvested at their logarithmic phase of growth and lysed by bead beating (30 s on/30 s off x 4 cycles). Equal amounts (∼50 µg) of proteins (quantitated using the BCA method) were subjected to SDS-PAGE analysis and transferred to PVDF membranes. Each blot was probed with specific antibodies. Proteins were detected using Enhanced Chemiluminescence (ECL, Thermo-Fisher).

### DIC and Fluorescence microscopy

To microscopically visualise cellular phenotypes, pellets from 1.5 mL of each culture were harvested and resuspended in 200 µL of PBS (*E. coli*) or 200 µL of 20% Tween 80 (*M. smegmatis*). For fluorescence microscopy the ORF corresponding to *MSMEG_3743* was cloned in pTetO-mRFP as an mRFP fusion protein with an intervening 5x Glycine linker, using the primers listed in Table S1. DAPI was used at a concentration of 2 ng/µl to stain bacterial nucleoids. DAPI and mRFP fluorescence was visualized using DAPI and DsRed channels respectively. Cells were passed through a syringe and 3-5 µL of these samples were loaded onto microscope slides harbouring a thin agarose (0.8-1%) pad followed by sealing with nail enamel. The cells were visualized at 100x under an oil immersion objective on either a Zeiss Axioimager Z1 or a Zeiss Axioplan 2 microscope. Cell numbers and cell lengths were measured manually.

### Gene Expression Analyses

To determine the levels of mRNA on *MSMEG_3743* overexpression, and for growth phase dependent gene expression studies, RNA was isolated from *M. smegmatis* mc^2^155/*M. tb* H37Ra using TRIzol reagent, as described by the manufacturer (Invitrogen). Following treatment with RNAse free DNAse I, cDNA synthesis was performed using the iScript cDNA synthesis kit (Bio-Rad) and subsequently used as a template for SYBR green based PCR amplification using gene specific primers designed to generate 200 bp amplicons (Table S1). To determine the growth phase dependent expression of *MSMEG_3743*, RNA was isolated from *M. smegmatis* mc^2^155 cultures harvested at OD of 0.7 (mid-log phase), 1.5 (late-log phase), 2.7 (early-stationary phase) and 3.7 (late stationary phase). Similarly, for the growth phase dependent expression analysis of *Rv1708*, RNA was isolated from cultures of *M. tb* H37Ra pelleted down at ODs of 0.8 (mid-log phase), 1.2 (early-log phase) and 1.5 (late-log phase). The level of each mRNA was normalized to the transcript levels of *M. smegmatis/M. tb sigA*. Relative fold changes were calculated with reference to transcript levels of each gene at the mid-log phase which was assigned a value of 1. At least two independent replicates were performed.

### Expression and purification of MSMEG_3743 and its site-directed mutants

To obtain protein for biochemical analyses, the ORF corresponding to *MSMEG_3743* was cloned into pET22b with a C-terminal 6x-His tag. The K52A and K57A mutations were generated by site directed mutagenesis using the primers in Table S1. For protein expression, the recombinant pET22b plasmids were transformed into *E. coli* BL21(DE3). Cultures were induced at 37°C for 4 h with 80µM IPTG. Recombinant proteins were purified by Ni-NTA affinity chromatography. Purification of MSMEG_3743 required an additional gel filtration step.

### ATPase Assays

ATP hydrolysis assays were performed as follows: 40-140 µM protein, 0.5 mM γ-^32^P- labelled (3000 Curie/mole) and 0.5 mM unlabelled ATP were mixed in 5x reaction buffer containing 100 mM HEPES-free acid, 25% glycerol, 250 mM potassium acetate, 0.5 mM DTT, 0.5 mg/ml BSA and 25 mM Magnesium chloride. The reaction was allowed to proceed for 30 min at 30°C. Equal reaction volumes of each sample were loaded on pre-run dried Ethyleneimine cellulose Thin Layer Chromatograms using 3.2% Ammonium bicarbonate solution as the mobile phase. After TLC, the gel was dried, and after a 10-30 min exposure, the radiograph was developed using a Phosphorimager. BSA and Thioredoxin were used as negative controls, while the S-190 extract from *Drosophila melanogaster* embryos was used as positive control. Suramin sodium was used as a non-specific ATPase inhibitor. All blots were subjected to densitometric scanning and the decrease in γ-^32^P-ATP spot was quantitated using ImageJ software.

### In silico analyses

*E. coli* DNA & protein sequences were obtained from the Colibri database (http://genolist.pasteur.fr/Colibri/), *M. tb* DNA & protein sequences were obtained from the Tuberculist database (http://tuberculist.epfl.ch/) [22]. *M. smegmatis* DNA & protein sequences for unannotated genes were obtained by performing BlastP analysis of the corresponding *M. tb* sequence using the JCVI-CMR database (http://blast.jcvi.org/cmr-blast/). Other mycobacterial sequences were obtained from Genolist (genolist.pasteur.fr/GenoList) [23] and the KEGG genome database [24]. Sequence similarities were determined using the BlastP algorithm (http://blast.ncbi.nlm.nih.gov/Blast.cgi). Multiple Sequence Alignments were performed using ClustalW omega (https://www.ebi.ac.uk/Tools/msa/clustalo/) [25, 26], and the output files were imported into ESPript 3 (https://espript.ibcp.fr)[27] to generate the formatted alignments (along with PDB 3Q9L as reference).

### Comparative Structural analysis of *E. coli* MinD and predicted structures of Rv1708 & Rv3660c

Structure coordinates for *E. coli* MinD were downloaded from the RCSB PDB database [28]. The predicted structure coordinates for Rv1708 and Rv3660c were downloaded from Alphafold [29]. Using the PDBsum server, it was confirmed that there were no Ramachandran outliers in the predicted structure [30]. The monomeric model of the Rv1708 was docked to obtain the dimeric Rv1708 model using the RosettaDock tool [31]. The structures were visualized, analyzed and compared using PyMOL.

### Statistics

For all experiments, the student’s T-test was conducted to determine statistical significance between two groups, when required.

## RESULTS

### Mycobacterial homologues of *Ec* MinD complement an *E. coli* Δ*minDE* strain to varying degrees

Since there were no reports of the Min and Nucleoid occlusion systems in Mycobacteria, we first performed pairwise sequence alignments of *E. coli* MinD with two putative pairs of Mycobacterial homologues of MinD [32]. The two candidate pairs, Rv1708/MSMEG_3743, Rv3660c/MSMEG_6171 show 23-25% sequence identity to *Ec* MinD, but show a much higher degree of homology among themselves (Fig. S1). To test if these proteins indeed encode MinD like functions, ORF’s corresponding to *Rv1708, MSMEG_3743, Rv3660c* and *MSMEG_617*1 were cloned into the *E. coli* expression vector pTrc99A, and transformed into *E. coli* HL1, a strain carrying a Δ*minDE* deletion. This strain shows a minicell phenotype and has earlier been used for the identification of the chloroplast encoded MinD in *Arabidopsis thaliana* [33]. In the complementation experiments (Fig. 1) both *MSMEG_3743* and *Rv1708* were observed to complement *E. coli* HL1 to levels comparable to *Ec minD*, while *MSMEG_6171* and *Rv3660c* showed a partial complementation phenotype. Neither *Ec minD* nor its mycobacterial homologues were able to complement *E. coli* RC1, a strain carrying a deletion in the entire *minB* (Δ*minCDE*) operon (data not shown). Our observations strongly indicate that Rv1708/ MSMEG_3743 are functional homologues of *Ec* MinD, and their high degree of conservation across mycobacteria (Fig. S1), points to their functional importance to mycobacterial physiology. This result led us to examine the properties of these homologues, relevant to their predicted function.

**Figure 1:**
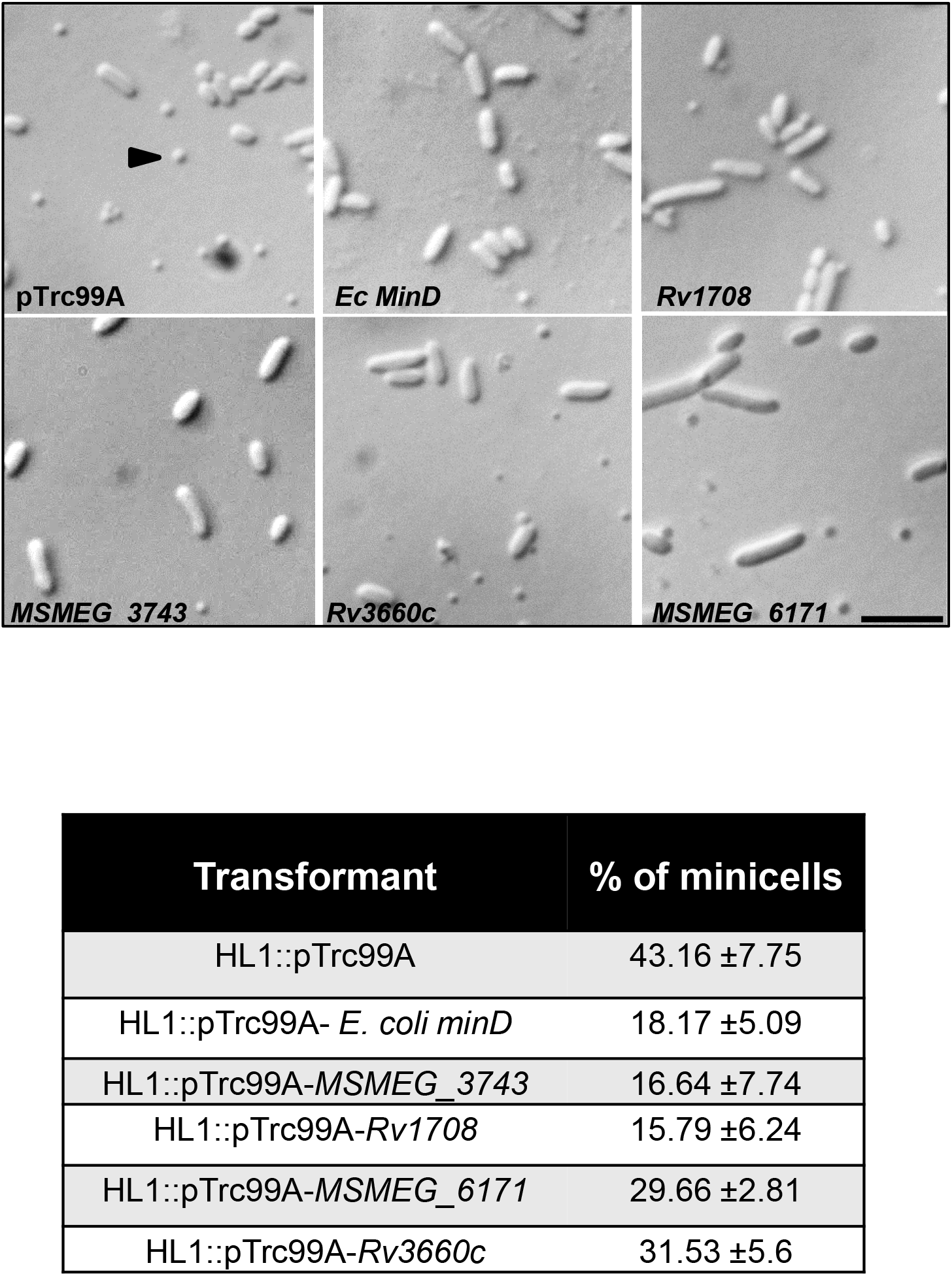
Representative DIC images *E. coli* HL1 transformed with *E. coli minD* and its predicted mycobacterial homologues. The arrowheads indicate the minicell phenotype; the scale bar is 5 µm. The table shows the frequency of minicells in these transformants; N=1000.

### Consequences of overexpressing *MSMEG_3743* and *MSMEG_6171* in *M. smegmatis*

Having identified *MSMEG_3743* as a fully complementing, and *MSMEG_6171* as partially complementing *M. smegmatis* homologues of *Ec minD,* we examined the consequences of overexpressing these genes in *M. smegmatis* using a Tet inducible system [34, 35]. In this experiment we assessed filamentation and viability (OD and CFU) following overexpression (Fig. 2). *MSMEG_3743* overexpression led to filamentation, a hallmark property of bona fide cell division genes. In addition, this led to a drastic change in CFU and significant change in OD (Figs 2c and 2d), clearly demonstrating the importance of its optimal expression. In contrast, overexpression of *MSMEG_6171*, neither caused *M. smegmatis* to filament, nor did it lead to changes in OD or CFUs (Figs 2c and 2d). This data is suggestive of MSMEG_3743 being a true homologue of *Ec* MinD, and MSMEG_6171 possibly being a member of the big MinD/ParA family. This filamentation phenotype was specific for *M. smegmatis*, since overexpression of *MSMEG_3743* did not cause any morphological changes, when it was overexpressed in *E. coli* MG1655 (data not shown). We observed a similar effect when *MSMEG_3743* was overexpressed using pJEX55 from the *hsp*60 promoter [36] (data not shown).

**Figure 2:**
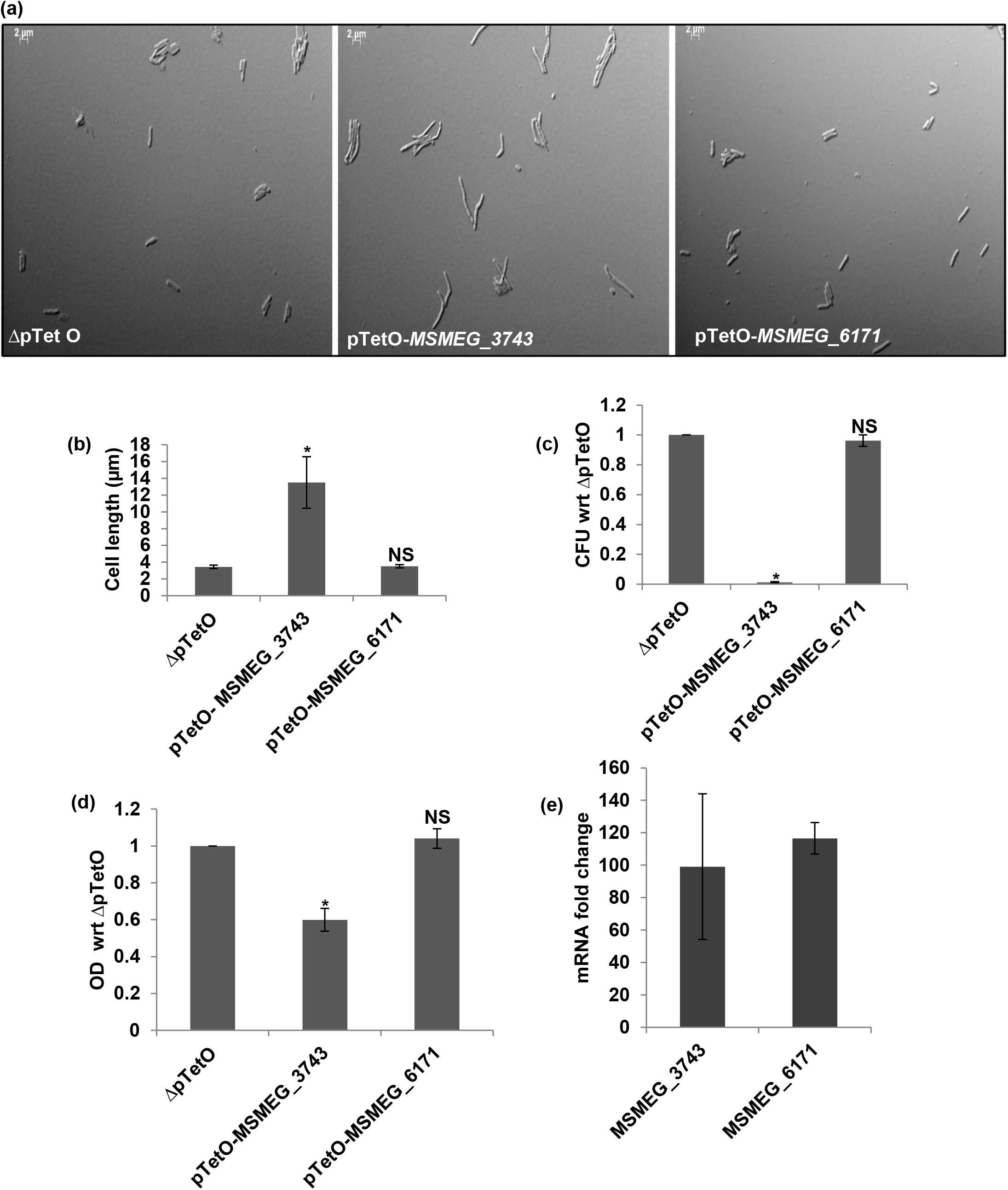
Effect of overexpression of *MSMEG_3743* & *MSMEG_6171* in *M. smegmatis*. (a) Representative DIC images of overexpression strains; Scale bar 2 µm (b) Average cell lengths of overexpression strains (µm) Error bars represent SEM. (c) CFU counts, represented as a ratio wrt ΔpTetO (d) OD measurements represented as a ratio wrt ΔpTetO (e) Transcript levels of *MSMEG_3743* & *MSMEG_6171* wrt ΔpTetO following overexpression, normalized to *Ms sigA.* *P<0.05, NS - not significant.

### ATPase activity of MSMEG_3743

The activity of *Ec* MinD is ATP dependent and *Ec* MinE has been shown to activate MinD *in vitro* [37, 38]. Like *Ec* MinD, MSMEG_3743 also has a conserved deviant Walker A motif associated with ATP hydrolysis activity, which led us to test its ATPase activity using a highly sensitive γ-^32^P-ATP based ATP hydrolysis assay. In this assay, the *Drosophila melanogaster* S-190 embryo extract was used as positive control due to the presence of multiple chromatin remodelers, which show ATP activity [39, 40]. BSA & Thioredoxin, which are not known to possess ATPase activity, served as negative controls for this assay. As shown in the radiograph in Fig. 3, a distinct decrease in the γ-^32^P-ATP band was observed in both the positive control as well as MSMEG_3743 lanes, revealing the ATP hydrolysis activity of this protein. We also tested the effect of the ATPase inhibitor Suramin [8-(3-Benzamido-4-methylbenzamaido)Napthalene-1,3,5-triSulphonic acid)] Sodium on this activity. The mode of enzyme inhibition (competitive/Non-competitive) by this compound has been observed to be enzyme specific [41]. We observed a dose dependent reduction in the amount of γ-^32^P-Pi released, clearly indicative of an inhibition of ATP hydrolysis (Fig. 3a). These results unequivocally establish that MSMEG_3743 possesses ATP hydrolysis activity. This observation however, does not rule out the possibility that this activity may be regulated by an activator as is characteristic of the ParA/MinD family of proteins.

**Figure 3:**
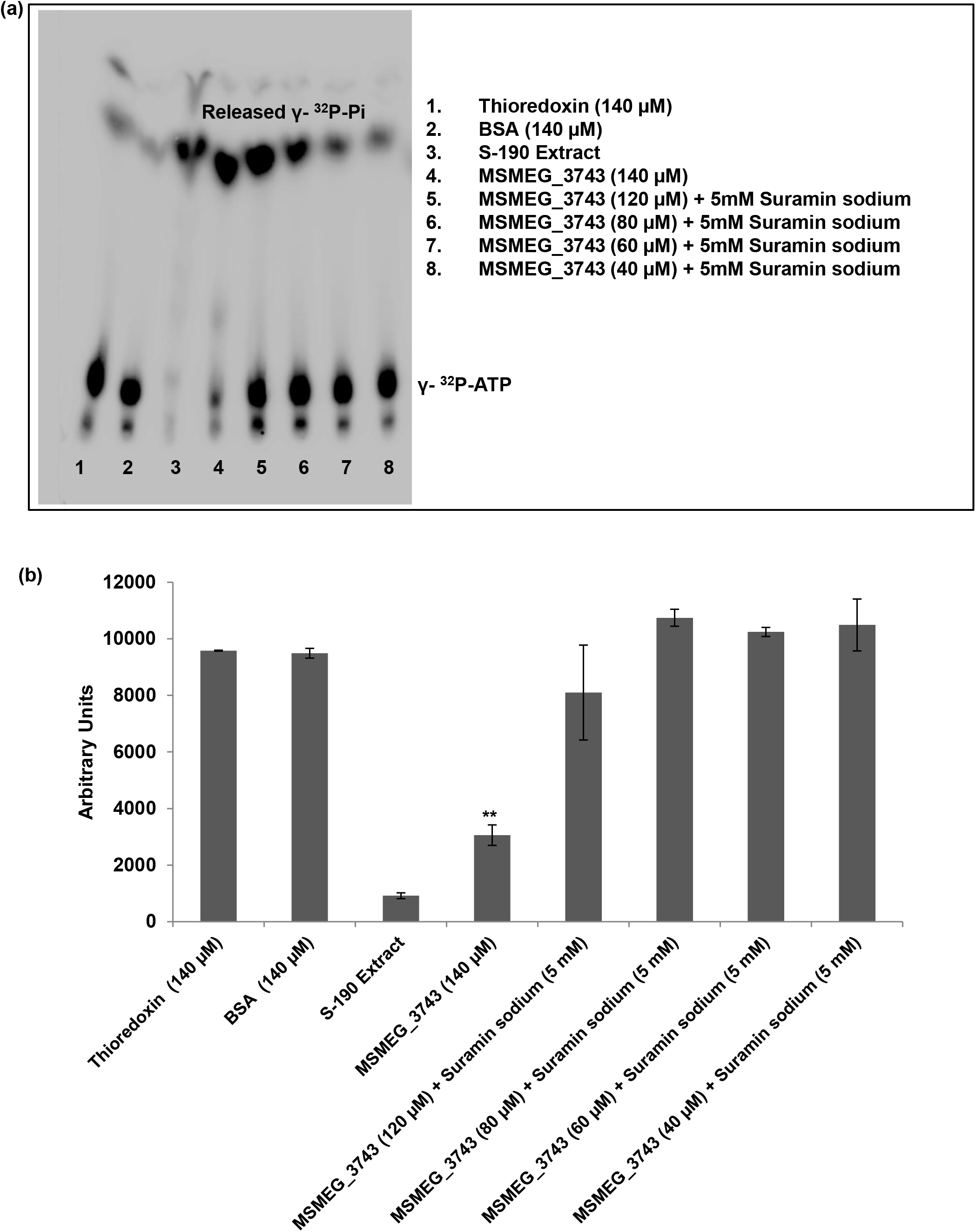
ATPase activity of MSMEG_3743. **(a)** Radiograph of γ^32^P-Pi release from hydrolysis of radiolabelled ATP. **(b)** Densitometric quantitation of unhydrolysed γ^32^P-ATP from each lane of the radiograph. Error bars represent SD; ** P<0.01, T-test done wrt BSA.

### Delineating the functional role of the Lysines in the Deviant Walker A motif of MSMEG_3743

As shown in Figure S2, there is high conservation of the Deviant Walker A motif in the MSMEG_3743/Rv1708 homologues across Mycobacteria. This motif differs from the Classical Walker A motif in the presence of a second lysine (referred as Signature lysine) other than conserved lysine present in the C-terminus of the motif. This signature lysine is known to be important for interaction with MinC, dimerization and ATPase activity [28, 42, 43]. The conserved lysine in the classical motif has also been shown to be involved in ATPase activity [44]. The currently accepted model suggests an interaction of the signature Lysine of *Ec* MinD with ATP bound to the other protomer [28, 43]. The rate of ATP hydrolysis increases 10 fold, when MinE binds to phospholipid-associated MinD, which leads to MinD-ATP dissociation from membrane bound MinE. To evaluate the functional importance of the two lysines at positions 52 and 57 in MSMEG_3743, we constructed site directed mutants, where the lysines were replaced with alanine residues and performed ATPase assays with the mutant proteins. We did not observe any significant change in the ATPase activity of the K52A and K57A mutants in comparison to wild type MSMEG_3743 (Figure S3) as observed in other members of the ParA/MinD family, where lysine substitution does not affect ATPase activity (*Eubacterium rectale* TadZ shows intrinsic ATPase activity despite the loss of both conserved lysines during the course of evolution)[45]. Since the conserved and signature lysines seemed to play no role in the ATPase activity of MSMEG_3743, we tested if these were involved in other functions as is known for members of the ParA/MinD family. The genes corresponding to K52A and K57A mutants of *MSMEG_3743* were overexpressed in *M. smegmatis* using a Tet inducible system along with the wild type gene and their ability to induce filamentation was assessed. Interestingly, overexpressing the K57A mutant was observed to induce filamentation to a significantly less degree as compared to WT *MSMEG_3743*, but did not affect the reduction in CFUs (Figure 4). Also, mutagenesis of the signature lysine did not seem to affect the function of MSMEG_3743 (Figure S4). These observations point to a role for K57 in the filamentation inducing property of MSMEG_3743, and suggest that this activity may be separable from its ability to cause a decrease in viability on overexpression.

**Figure 4:**
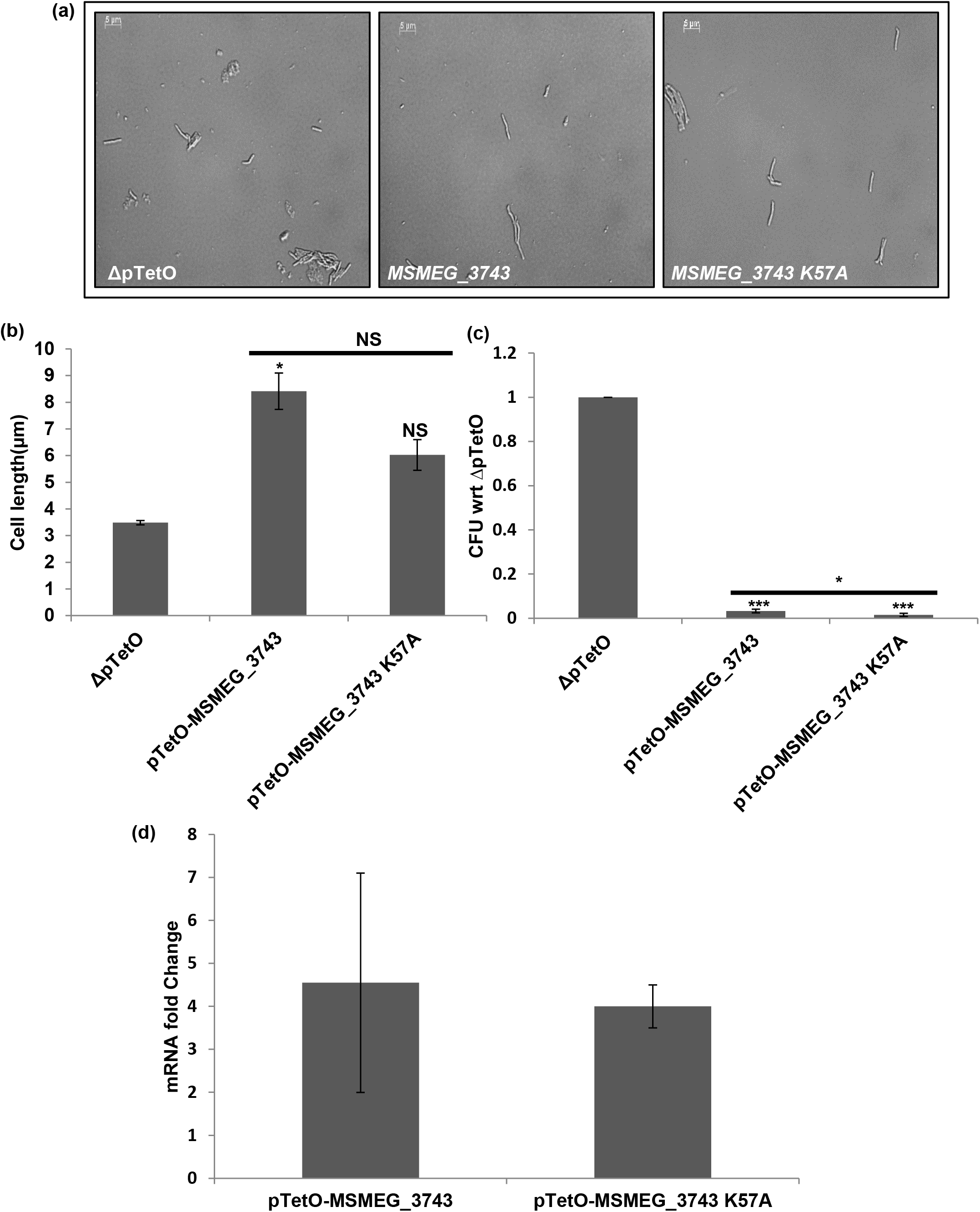
Effect of overexpression of *MSMEG_3743 K57A* in *M. smegmatis*. (a) Representative DIC images of overexpression strains; Scale bar 2 µm (b) Average cell lengths of overexpression strains (µm) Error bars represent SEM. (c) CFU counts, represented as a ratio wrt ΔpTetO (d) Transcript levels of *MSMEG_3743* & *MSMEG_3743 K57A* following overexpression, wrt *Ms sigA.* ** P<0.01, * P<0.05, T-test performed wrt ΔpTetO; NS – Not Significant.

### Cellular localization of MSMEG_3743

Members of the ParA/MinD/SojA family are known to localize to different parts of the cell like the mid cell, or the pole, to perform their various functions. In order to determine its localization pattern, the ORF corresponding to *MSMEG_3743* was cloned in a Tet inducible vector as both N- and C-terminal translational fusions with mRFP (modified RFP) with a 5x glycine linker between the these two regions. To test, if the addition of the mRFP tag affect the function of MSMEG_3743, both fusion constructs were tested for their ability to induce filamentation and reduce viability on overexpression in *M. smegmatis*. Only the C-terminal fusion construct was observed to behave like the wild type protein (data not shown) and was therefore retained for further analyses. On its overexpression MSMEG_3743-mRFP showed longitudinal equidistant localization (Figure 5a) as has been reported for PpfA (*Rhodobacter sphaeroides*)[46], ParA (*Synechococcus elongatus*)[47], ParA in the P1 partition system [48] and for Rv1708 [49], the *M.tb* homologue of MSMEG_3743. Since, overexpression of this construct with 50 ng/ml ATc induction led to filamentation, which does not represent the physiological state of bacteria, we performed a range finding experiment for ATc to identify a concentration under which there were no significant changes in the cell length of recombinant *M. smegmatis* expressing this construct but a sufficient level of mRFP expression for the visualization of protein localisation. We observed that at concentration of 1 ng/ml, both these conditions were satisfied and used this induction condition to visualise MSMEG_3743. The protein showed both polar and apolar localisation (Figure 5b), and appears to move from the newly formed pole to the mid-cell during the cell cycle.

**Figure 5:**
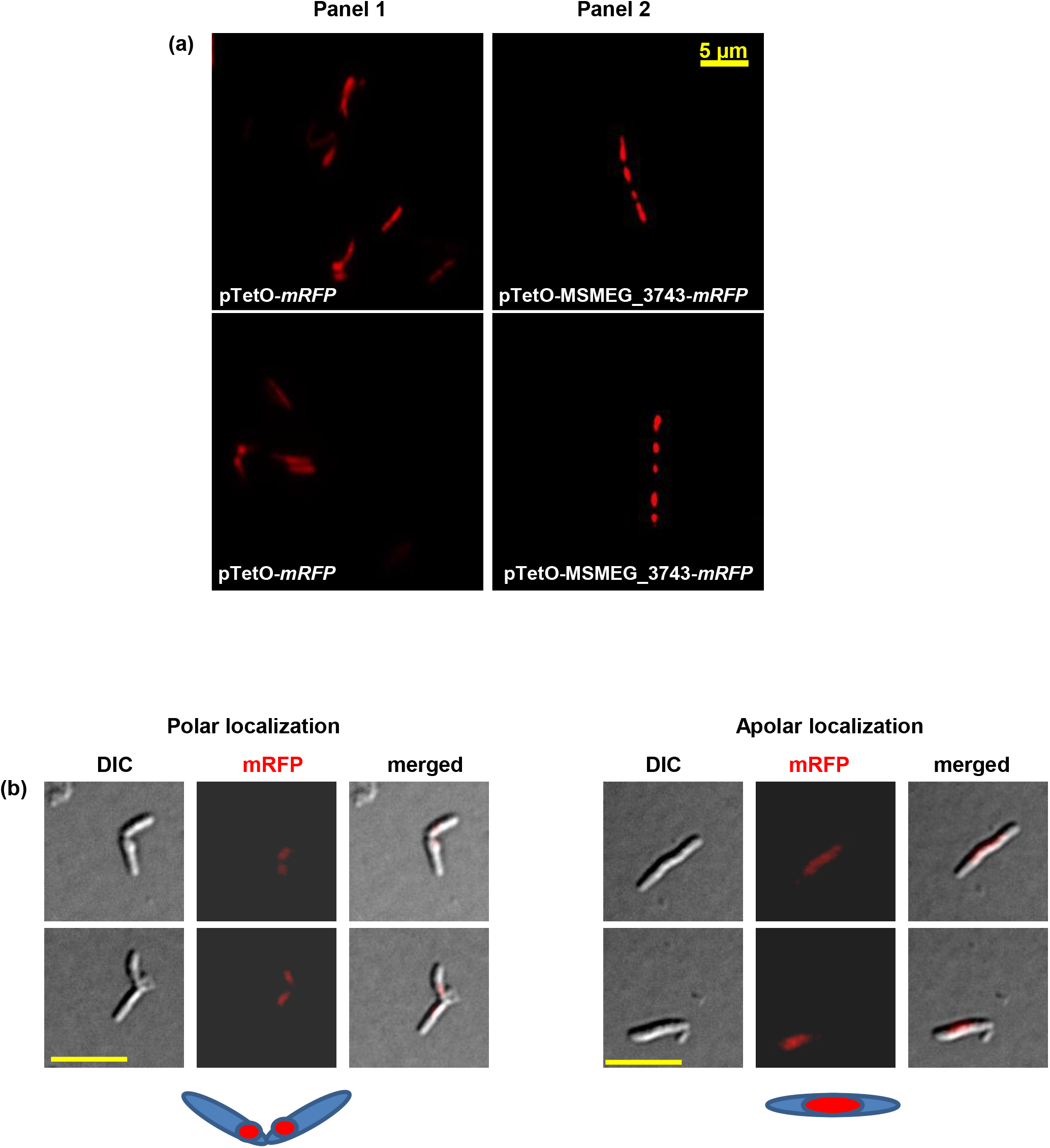
Localization of MSMEG_3743-mRFP fusions in *M. smegmatis*. (a) Representative images showing localization of MSMEG_3743-mRFP under conditions of overexpression (50 ng/ml ATc); pTetO-mRFP represents the control. (b) Representative images and schematic showing polar and apolar localization of MSMEG_3743-mRFP under conditions of native expression (1 ng/ml ATc); Scale bar, 5 µm.

### Subcellular localization of MSMEG_3743

Most divisome proteins are either localized to either the cytoplasm or are membrane bound. In order to determine its subcellular localization we used an Acetamide inducible system to express *MSMEG_3743* as a His-tagged fusion protein in *M. smegmatis*, and prepared its subcellular fractions. These fractions were resolved on SDS-PAGE and subjected to western blotting using an anti His-tag antibody. MSMEG_3743 was observed to predominantly localize to the cell wall and cell membrane fractions (Figure 6). This opens up the possibility of other membrane localized divisome proteins to interact with MSMEG_3743 and indirectly affect the process of FtsZ ring assembly.

**Figure 6:**
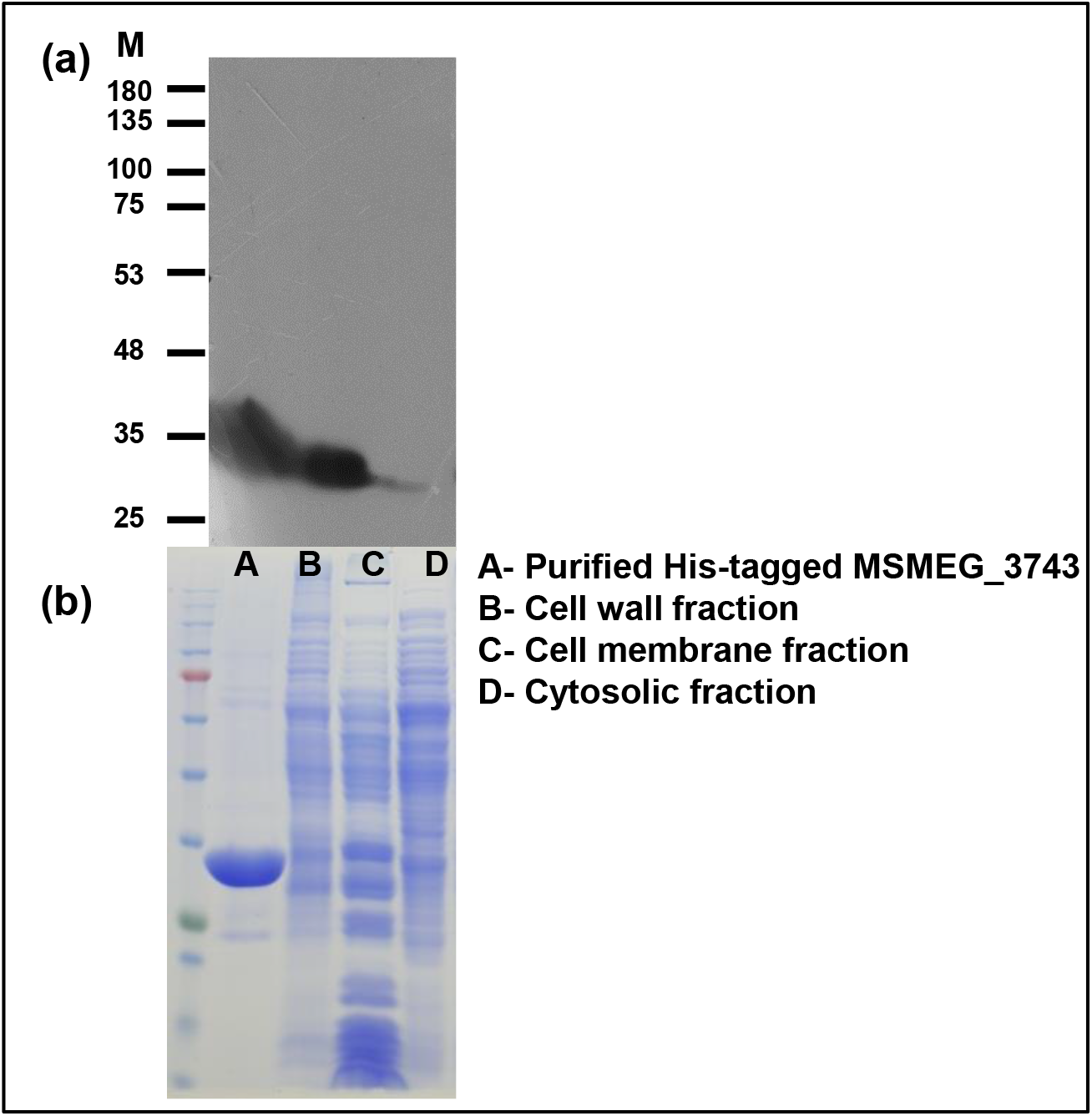
Sub-cellular localization of MSMEG_3743. (a) Western blot of sub-cellular fractions of *M. smegmatis* expressing His-tagged MSMEG_3743 probed with an anti His-antibody (b) Coomassie stained gel showing equal loading of all fractions; M -Size marker, KDa.

### Identification of interacting partners of Rv1708/MSMEG_3743

Protein-protein interaction is crucial to divisome formation since this involves the assembly of proteins/protein complexes in defined stoichiometry [50]. We hypothesised that since Rv1708/MSMEG_3743 are involved in divisome formation, they must interact with other proteins while performing their function. To identify their interacting partners, candidate proteins predicted to interact with Rv1708 were chosen [20] and their interactions tested using Mycobacterial Protein Fragment Complementation (MPFC)[21]. In the assay, co-transformants of *M. smegmatis* containing *Rv1708* with *M.tb scpA* or *M.tb parB* showed growth on 20 µg/ml Trimethoprim containing plates, indicating that both these chromosome segregation proteins interact with Rv1708 (Figure 7a) Both these interactions were validated biochemically, using pull-down assays (Figure 7b and 7c). Extrapolating these observations, we were able to establish using MPFC that MSMEG_3743 interacts with MSMEG_6938, the *M. smegmatis* homologue of *M.tb* ParB (Figure 7d), implying that this interaction is functionally important for mycobacterial cell-division.

**Figure 7:**
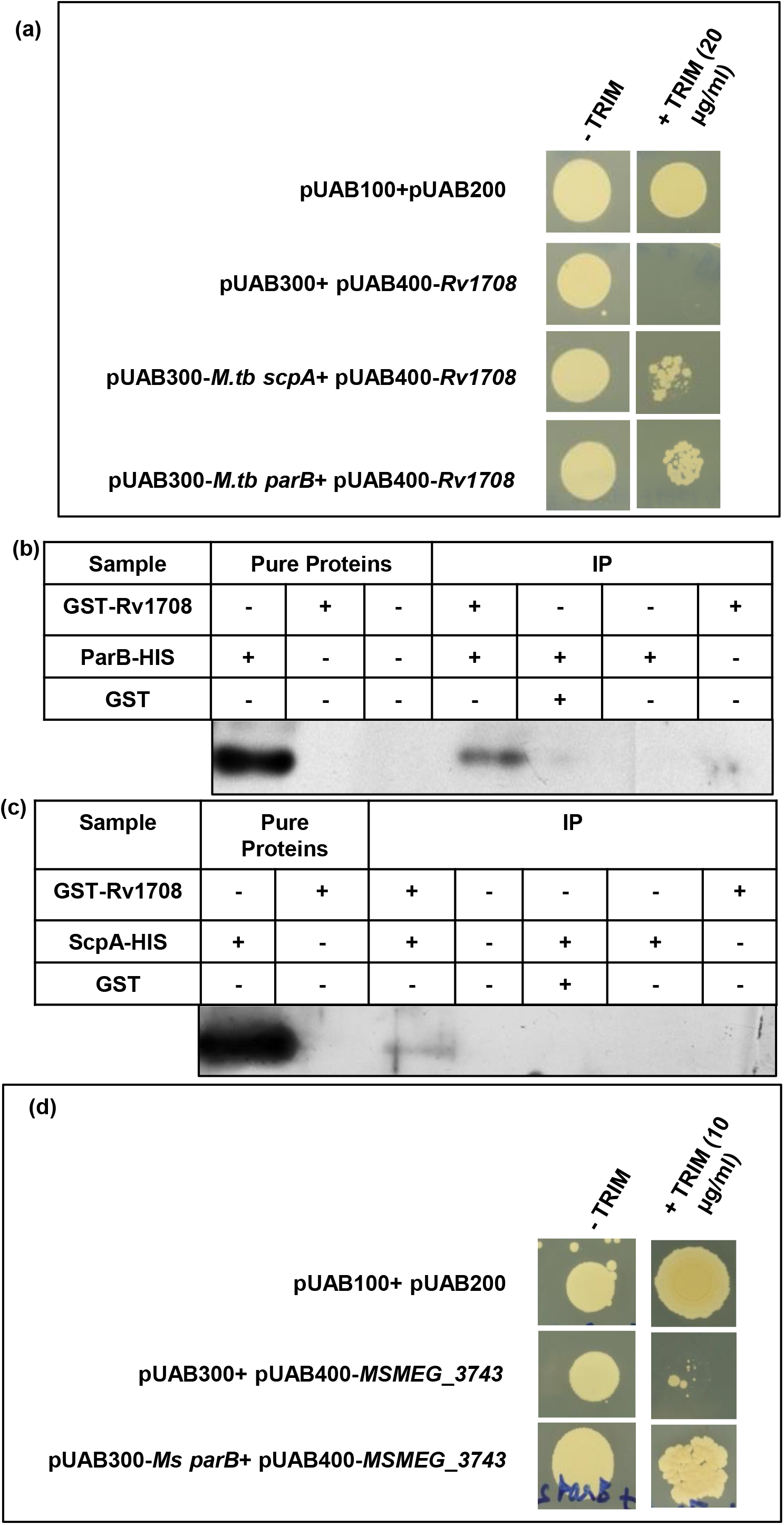
Identification of Interacting partners for Rv1708/MSMEG_3743. (a) MPFC showing interaction of Rv1708 with *M.tb* ParB and *M.tb* ScpA. (b, c) Biochemical validation of Rv1708-*M.tb* ParB and Rv1708-*M.tb* ScpA interactions respectively.(d) MPFC showing interaction of MSMEG_3743 with *Ms* ParB.

### Effect of *Ms parB* overexpression in *M. smegmatis*

In order to assess the functional role of the observed interactions, *Ms parB* (*MSMEG_6938*), was over expressed in *M. smegmatis* using the Tet inducible system described previously. In addition we also overexpressed *Ms scpA* (*MSMEG_3742*), since it was observed to interact with Rv1708, the *M.tb* homologue of MSMEG_3743. As shown in Figure 8, overexpression of *Ms parB* phenocopied the effects of *MSMEG_3743* overexpression and led to both filamentation of *M. smegmatis* as well as a drastic reduction in viability. This strongly suggests that *Ms ParB* and *MSMEG_3743* act in the same pathway. ParB is activator of the ATPase activity of ParA activity and helps in recruiting SMC and the replisome [51, 52]. Since overexpression of both *MSMEG_3743* and *Ms parB* leads to filamentation, this shows direct role for these proteins in mycobacterial cell division. The mechanism by which these proteins mediate this effect is currently unclear to us.

**Figure 8:**
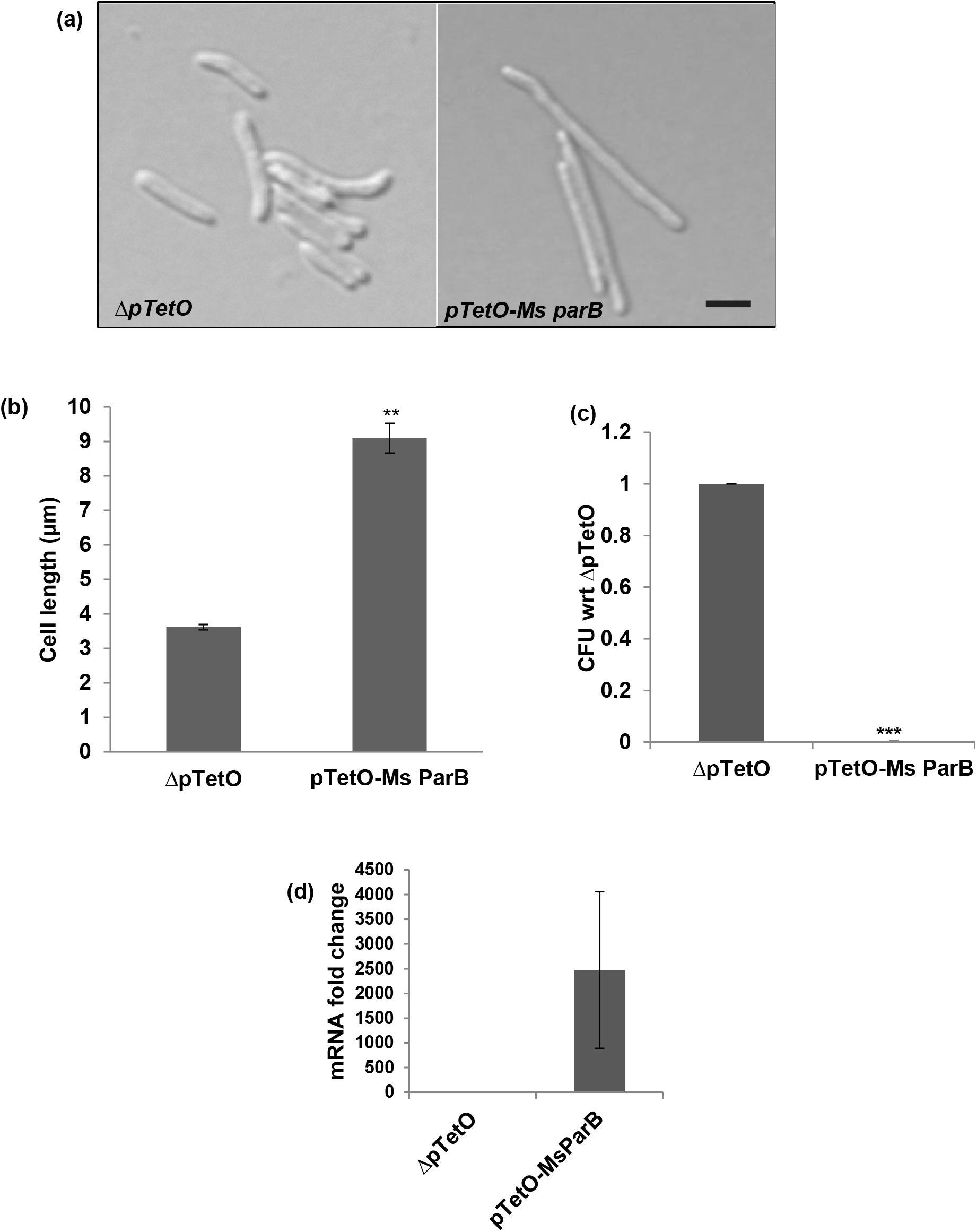
Effect of overexpression of *Ms parB* (*MSMEG_6938*) *in M. smegmatis*. (a) Representative DIC images of overexpression strains; Scale bar 2 µm (b) Average cell lengths of overexpression strains (µm) (c) CFU counts, represented as a ratio wrt ΔpTetO (d) Transcript levels of *Ms parB* following overexpression, wrt *Ms sigA.* Error bars represent SEM. *** P<0.001, ** P<0.01, T-test performed wrt ΔpTetO.

### Interaction of MSMEG_3743 with the bacterial chromosome

*Ec* MinD has been shown to bind chromosomal DNA and a mutation at Arginine 219 leads to an aberration in chromosome segregation [53]. To test if MSMEG_3743 shares this same property, we stained *M. smegmatis* cells expressing an *MSMEG_3743-mRFP* fusion with DAPI and performed fluorescence microscopy. We observed co-localisation of DAPI and mRFP fluorescence indicating that MSMEG_3743 was associated with the chromosome (Figure S5). Since MSMEG_3743 is mostly localized to the cell wall and partially to cell membrane, this association might be indirect. Based on our protein-protein interaction study, it would not be incorrect to hypothesise that this interaction may be mediated by ParB, which is known to bind to proximal *parS* sequences on the *M. smegmatis* chromosome. Out of the 5 suggested *parS* sites, 3 have been shown to bind with *Ms* ParB. Two of these are proximal to the ori-site, which further recruits ParB by ParB-ParB interactions and spreads to the nearby regions on the chromosome [54].

### Gene expression analyses of *MSMEG_3743/ Rv1708* during different phases of growth

In order to determine the gene expression patterns of *MSMEG_3743* and *Rv1708* as a function of growth, RNA was isolated from *M. smegmatis* and *M.tb* at different growth phases, and their expression was determined using real-time RT-PCR. Expression of both genes was observed to decrease after log phase (Figure S6). It is likely that the activities of *MSMEG_3743* and *Rv1708* are optimally required only during phases of active growth and their expression is regulated by a growth phase sensor.

### Comparative Structural analysis of *E.coli* MinD and its putative *M.tb* homologues

To further validate our identification of Rv1708 as being a homologue of *E.coli* MinD, we performed comparative structural analyses to identify possible conserved structural elements in these two proteins. Alignment of selected sequences revealed an overall low relative sequence identity, but the deviant walker A motif was found to be conserved among all the proteins selected for the alignment (Figure S7). However, on structural analysis we were able to identify subtle features unique to both Rv1708 and Rv3660c in comparison to *Ec* MinD. The alignment in Figure 9a clearly shows that the overall architecture of *Ec* MinD and Rv1708 is similar, and their secondary structural elements overlap - the relatively high RMSD value of. 2.1 (Cα backbone) can be attributed to the low sequence identity between the two proteins. The ATP binding site and the orientation of the ATP molecule in Rv1708 coincides with that of *Ec* MinD, as does the geometry of the Walker A motif (Figure 9b). Additionally, we noticed a helix (sequence ^152^IDLSAAEIQLVN^163^) inserted between β4 and α5 of Rv1708 which appears to be disordered in *Ec* MinD (Figure 9d). Contrary to what we observed for Rv1708 only a part of Rv3660c modestly matches in structure with *Ec* MinD with an RMSD value of 3.2. Its N-terminal region spanning 1 to 112 amino acids contains additional structural elements, which are absent in *Ec* MinD (Figure 9c). It is probable that this region, which perfectly matches with the NTD of MSMEG_6171could be associated with a function distinct from *Ec* MinD. These findings suggest that although the Walker A motif is present in Rv3660c and MSMEG 6171, the true mycobacterial homologues of *Ec* MinD are Rv1708 and MSMEG 3743.

**Figure 9:**
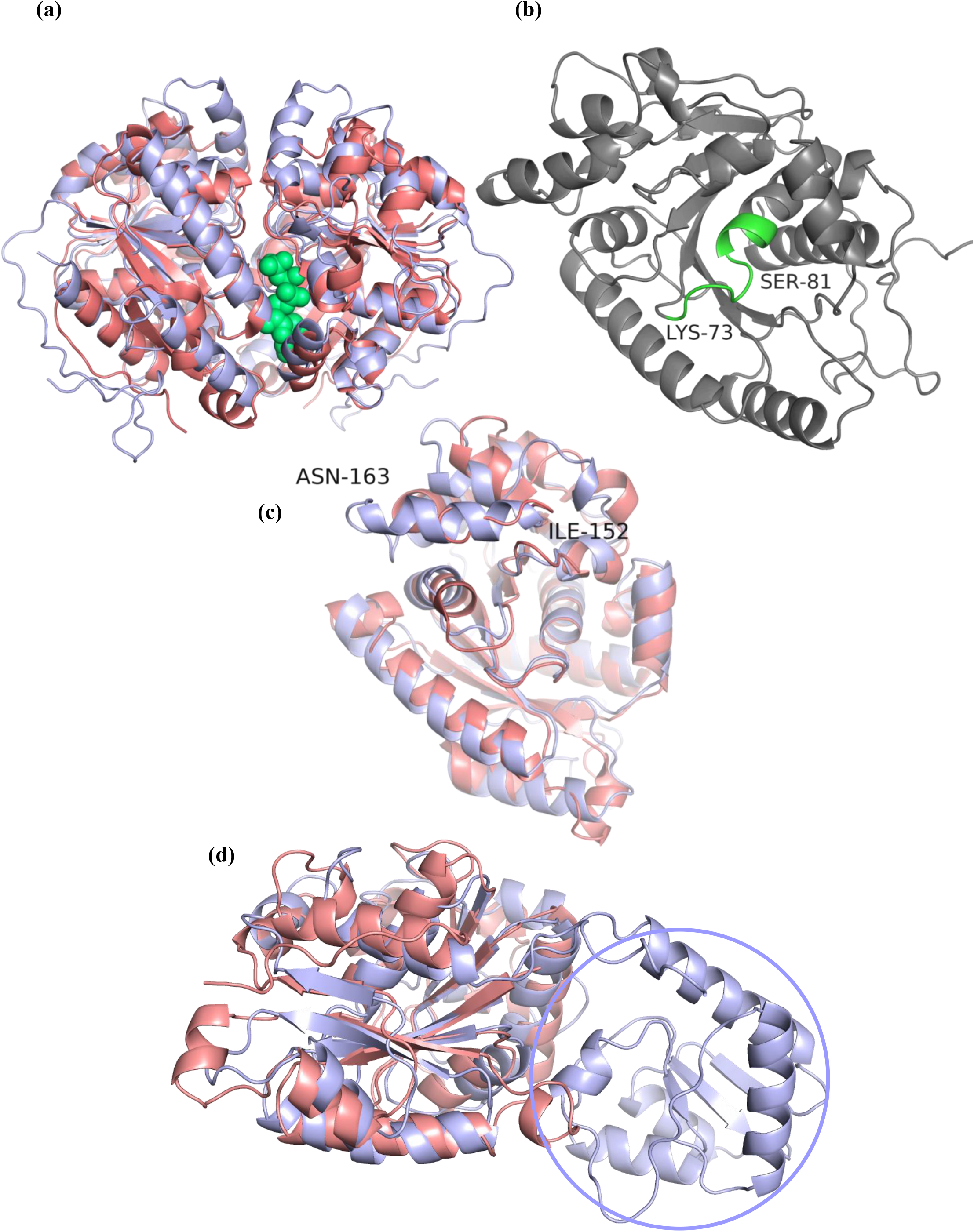
Structure based comparison of *E.coli* MinD and its putative *M.tb* homologues. (a) Structural alignment of Rv1708 (light blue) with *E. coli* MinD (pink). The ATP molecule (green) can be seen bound in the active site, suggesting conserved architecture of the active site in *E. coli* MinD and Rv1708. In both proteins, the ATP binding site is situated in the dimer interface. (b) General architecture of Rv1708 with the deviant Walker A motif highlighted in green. (c) Insertion of helix I152 to N163 observed in Rv1708 (light blue) which is disordered in *E. coli* MinD (pink) (d) Structural alignment of *E. coli* MinD (pink) with Rv3660c (blue) showing the absence of significantly shared structural elements between the two. The N-terminal region encircled in Rv3660c is absent in *E. coli* MinD.

## DISCUSSION

The adaptability of *Mycobacterium tuberculosis* in being able enter into a dormant state under hostile conditions and undergo resuscitation, when the environment is favourable is one of the reasons for its pathogenic success. Since metabolic shutdown during dormancy renders *M.tb* resistant to drug treatment, it is vital that we understand the mechanism of mycobacterial cell division and the players involved in this complex process. In order to identify mycobacterial homologues of *Ec* MinD, we used a target based complementation approach using the *E. coli* HL1 mutant (Δ*minDE*), and showed Rv1708/MSMEG_3743 to be true homologues of *Ec* MinD. *Rv3660c* and *MSMEG_6171*showed a partial complementation phenotype, as did *Rv3213c/ MSMEG_1927,* another predicted mycobacterial homologue of *Ec* MinD [55] (data not shown). The comparative structural analysis of predicted structures of Rv1708 and Rv3660c with the crystal structure of ATP bound *Ec* MinD dimer unequivocally showed that Rv1708 overlaps better with *Ec* MinD in comparison to Rv3660c, providing validation to our complementation results.

*MSMEG_3743* was observed to lead to filamentation when overexpressed in *M. smegmatis*, a hallmark of proteins involved in formation of the septum. From the drastic decrease in CFUs on overexpression, we hypothesize that *MSMEG_3743* is an essential gene, and generation of a knock-out strain in all probability will require an episomal rescue of the gene.

Earlier studies had examined the ATP binding ability of mycobacterial proteins using either click chemistry based high throughput screening [56] or a chemical proteomics approach using Desthiobiotin-conjugated ATP as a molecular probe [57]. However, neither of these studies reported Rv1708/MSMEG_3743 to be an ATP binding/ATP hydrolyzing protein, possibly due to low levels of ATPase activity of Rv1708/MSMEG_3743, or due to the requirement of an activator, as shown for *Ec* MinD. Targets of ParA/MinD/SojA family members are often either DNA or the membrane [58] and their ATPase activity is activated by an activator (e.g. MinE), but members of the Orphan ParA subfamily [45, 59–64] are not associated with these usual partners. Some members like MipZ [64] and PldP [59] are involved in spatial regulation of Z-ring positioning in bacteria lacking MinD. Our detection of ATPase activity in MSMEG_3743 was consistent with the presence of a conserved Walker A motif, and site directed mutagenesis allowed us to identify a role for the conserved lysine, K57 in the filamentation inducing property of *MSMEG_3743*. Site directed mutagenesis of both Lysines in the Deviant A Walker Motif did not abolish the ATPase activity of MSMEG_3743, similar to *E. rectale* TadZ, where the function of the signature P-loop lysine is compensated for by a lysine residue from an adjacent α-helix [45].. Our data suggests that this activity may be distinct from its viability reduction property when overexpressed. It is conceivable the decrease in CFU may be due to the bacteriostatic effect of the expression of a MazF like toxin, as has been observed in *M.tb* [65].

In protein-protein interaction assays, *M.tb* ScpA and ParB were identified as binding partners for Rv1708, and the *M. smegmatis* homolog of *M.tb* ParB was observed to interact with MSMEG_3743.This observation is consistent with the tenet, that bacterial cell division is mediated by the concerted effect of protein complexes ScpA and ScpB that form the SMC (Structural Maintenance of Chromosome) Complex with the SMC protein scaffold, which helps in condensation and segregation of the bacterial chromosome. ParB is primarily localised to the cytoplasm and activates ParA to enable chromosome segregation - a knock-out of *parB* has been shown to cause defects in chromosome segregation in *Bacillus subtilis* [66], *Pseudomonas aeruginosa* [67], *Myxococcus xanthus* [68] and *Mycobacterium smegmatis* [69]. ParB is essential for septum site determination in *Caulobacter crescentus* [70] and *Corynebacterium glutamicum* [71]. This protein binds near the Ori region at two ParS DNA sequences like the ParA/MinD like protein MipZ of *C. crescentus*, which acts as a negative regulator of FtsZ polymerization. In mycobacteria, ParB has been shown to interact with ParA and Wag31 the homolog of *B. subtilis* DivIVA [69]. ParB plays an important role in the localization of replisomes near the mid cell and recruits the SMC protein near to the Ori site. Wag 31 localizes to the cell pole [72], since it has the intrinsic property of recognizing the most curved part of bacteria. *M.tb* [49], *C. glutamicum* [59], *M. Xanthus* [73], *C. crescentus* [74], *V. cholera* [75], *P. aeruginosa* [76], and *S. pneumonia* [77] share a similar Ori-Ter chromosome organization unlike that in *E. coli* [78]. In all these organisms, ParB plays an important role in chromosome organization and septum site determination. Caulobacter and Mycobacteria share similar properties like the absence of a Nucleoid Occlusion System, Ori-Ter Chromosome organization, essentiality of ParB in cell division & the role of a polar anchor (which plays a crucial role in ParAB and Ori region recruitment in bacteria containing the Ori-Ter Chromosome organization). *Ms* ParB has shown to be involved in asymmetric segregation of chromosomes during mycobacterial cell cycle [79].

Based on our findings, we propose that MSMEG_3743 participates in regulation of divisome formation along with other unknown proteins (Figure 10). While the mechanistic details of MSMEG_3743 function in septum formation remain to be elucidated, it is conceivable that this protein may function like the other ParA/SojA family members - PomZ of *M. xanthus* which marks the FtsZ polymerization site at mid cell, or like MipZ, a spatial regulator co-ordinating chromosome segregation with cell division in *C. crescentus*. Our data clearly establishes *Rv 1708* and *MSMEG_3743* as bonafide mycobacterial homologues of *E. coli MinD,* implying a role for a Min-like system in mycobacterial divisome formation. We propose that *Rv 1708* and *MSMEG_3743* be henceforth referred to as *M. tb minD* and *Ms minD* respectively.

**Figure 10:**
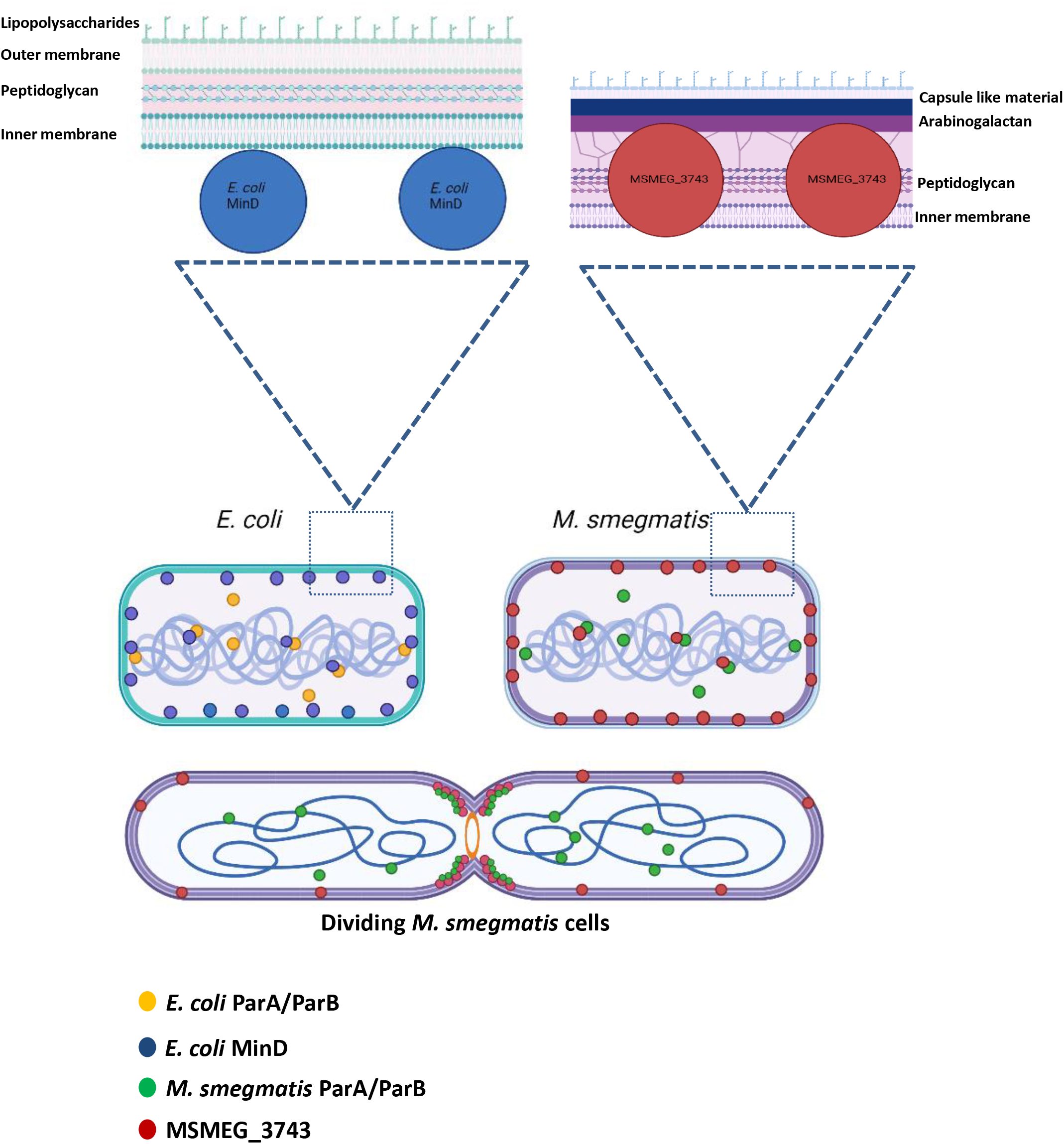
Model depicting the role of MSMEG_3743 in mycobacterial cell division. The Upper panel depicts the presence of ParA/ParB and MinD in both *E. coli* and *M. smegmatis* cells. *E. coli* MinD is present both in the cytoplasm and bound to the membrane, its interaction with the chromosome has also been demonstrated. Our data suggests MSMEG_3743 to be a true homologue of *E. coli* MinD, with subcellular localisation data showing the presence of MSMEG_3743 in the cell wall and cell membrane. Fluorescence microscopy data suggests co-localization of MSMEG_3743 with its chromosome. The Lower panel shows a dividing *M. smegmatis* cell forming two daughter cells, with MSMEG_3743 localised primarily at the septum (future new poles of daughter cells). The insets for the upper panel show the localisation of *E.coli* MinD and MSMEG_3743 in *E. coli* and *M. smegmatis* respectively, in the context of their cell envelopes.

## Acknowledgements

We gratefully acknowledge the help of Dr. Ameya Kumar Bendre, Department of Chemistry, Savitribai Phule Pune University, Pune India, in performing the comparative structural analysis of *E.coli* MinD and its putative *M. tb* homologues. We thank Dr. Swathi Chodisetty, National Centre for Cell Science, Pune, India for assisting in preparation of the model for MSMEG_3743 function.

## Author contributions

T.R.R. and V.K. designed the study. V.K. and S.S.G.S. performed the experiments.

T.R.R. and V.K.. analysed the data. T.R.R. and V.K. wrote the manuscript.

