## Supplementary Figures and Table for "Septum site placement in *Mycobacteria* - Identification and Characterization of mycobacterial homologues of *Escherichia coli* MinD"

| Subject / Query | *Ec* MinD (270 aa) | Rv1708 (318 aa) | Rv3660c (350 aa) | MSMEG_3743 (297 aa) | MSMEG_6171 (413 aa) |
|---|---|---|---|---|---|
| ***Ec* MinD (270 aa)** | 100 | Aligned 2-195 Identity 24% (50/212) Positives 48 % (103/212) | Aligned 4-150 Identity 23% (35/149) Positives 41 % (62/149) | Aligned 2-195 Identity 25% (52/212) Positives 48 % (102/212) | Aligned 4-150 Identity 23% (34/147) Positives 39 % (58/147) |
| **Rv1708 (318 aa)** | Aligned 64-261 Identity 24% (50/212) Positives 48 % (103/212) | 100 | Aligned 66-103 Identity 34% (13/38) Positives 52 % (20/38) | Aligned 21-318 Identity 85% (253/298) Positives 91% (272/298) | Aligned 66-103 Identity 39% (15//38) Positives 47% (18/38) |
| **Rv3660c (350 aa)** | Aligned 117-255 Identity 23% (33/146) Positives 41 % (61/146) | Aligned 267-342 Identity 25% (22/88) Positives 40 % (35/88) | 100 | Aligned 117-150 Identity 37% (14/38) Positives 55% (21/38) | Aligned 1-138 Identity 60% (203/338) Positives 71% (241/338) |
| **MSMEG_3743 (297 aa)** | Aligned 43-240 Identity 25% (52/212) Positives 48 % (102/146) | Aligned 2-297 Identity 81% (253/298) Positives 91 % (272/298) | Aligned 45-238 Identity 26% (23/207) Positives 36 % (75/207) | 100 | Aligned 40-82 Identity 40% (17/43) Positives 48% (21/43) |
| **MSMEG_6171 (413 aa)** | Aligned 183-296 Identity 23% (27/119) Positives 40% (48/119) | Aligned 183-216 Identity 39% (15/38) Positives 47 % (18/38) | Aligned 72-404 Identity 60% (203/338) Positives 71% (241/338) | Aligned 183-216 Identity 39% (15/38) Positives 50% (19/38) | 100 |

**Figure S1**: **Checkerboard showing sequence conservation of *E. coli* MinD and its mycobacterial homologues Rv1708/MSMEG_3743 and Rv3660c/MSMEG_6171**

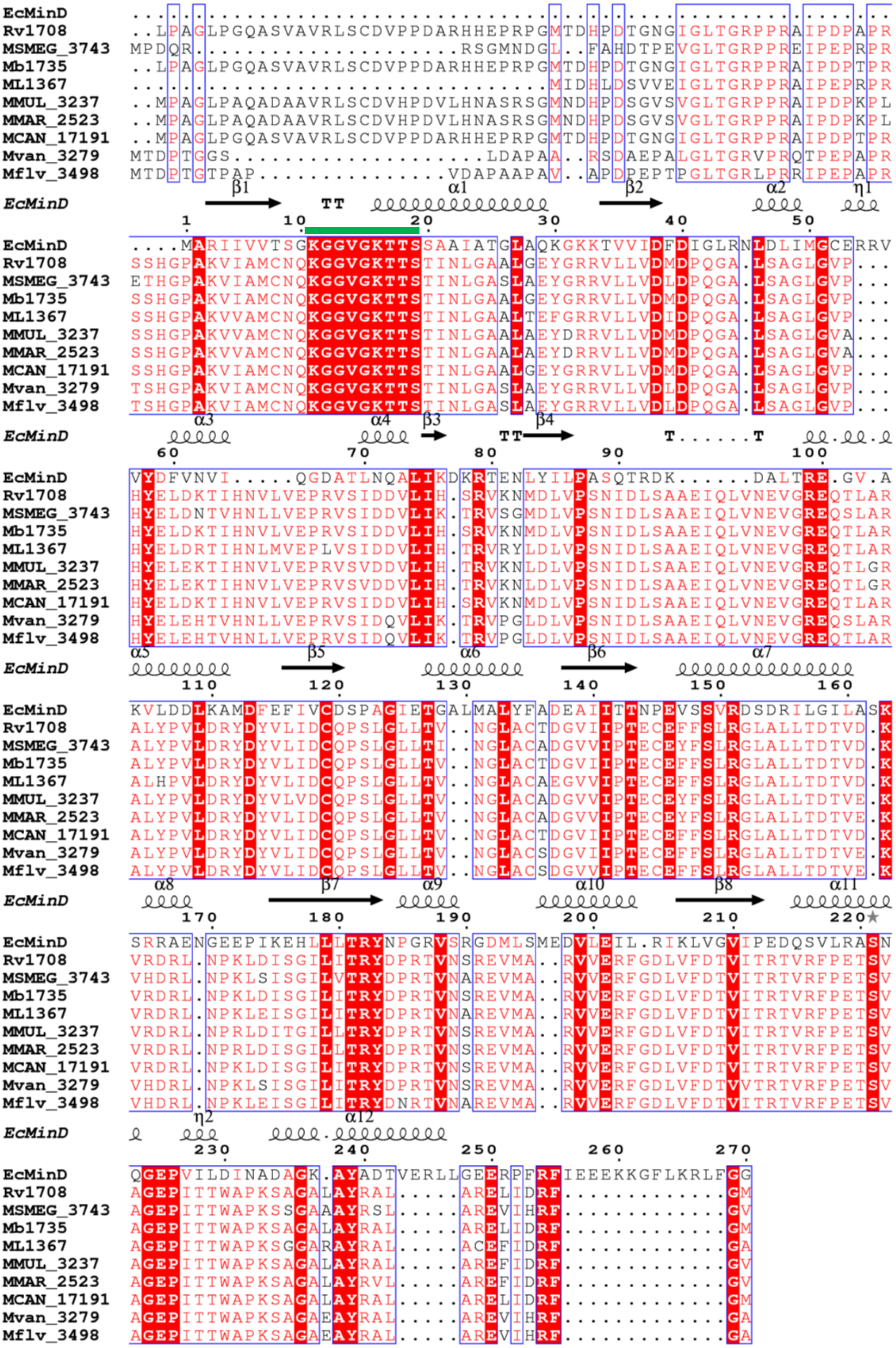

**Figure S2**: **Conservation of MSMEG_3743/Rv1708 sequences across mycobacteria.** The green line shows conservation of the Deviant Walker A motif. Mmul-*Mycobacterium ulcerans,* Mmar-*Mycobacterium marinum,* Ml-*Mycobacterium leprae,* Mcan-*Mycobacterium canetti,* Rv-*Mycobacterium tuberculosis,* Mb-*Mycobacterium bovis,* MSMEG-*Mycobacterium smegmatis,* Mvan-*Mycobacterium vanbaalenii,* Mflv-*Mycobacterium gilvum*.

(a)

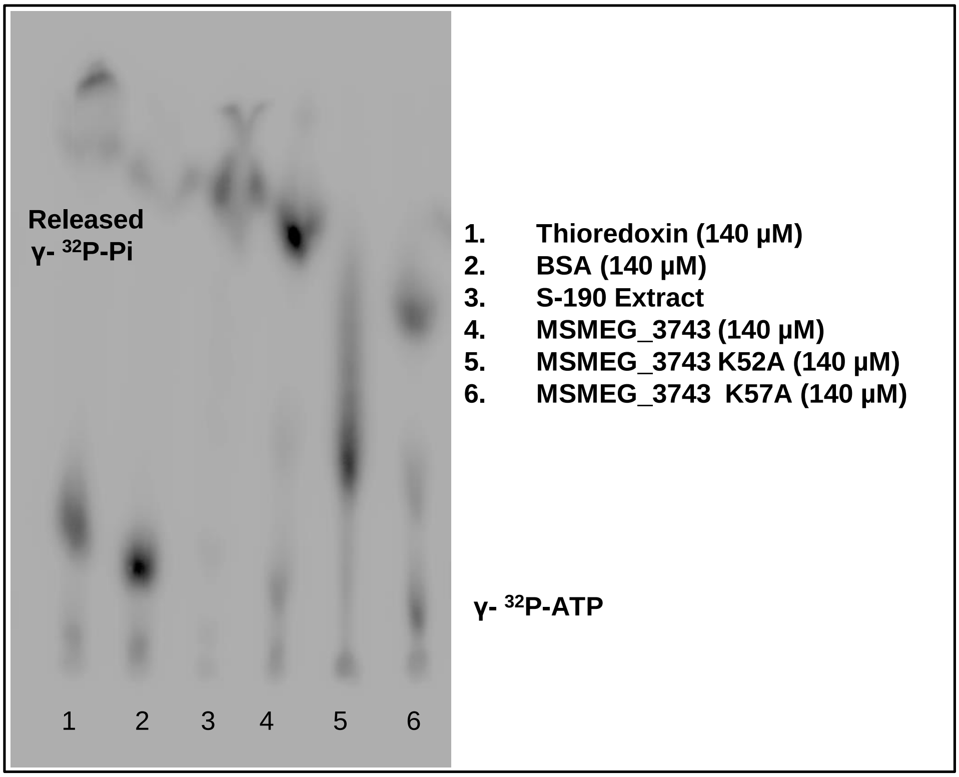


(b)

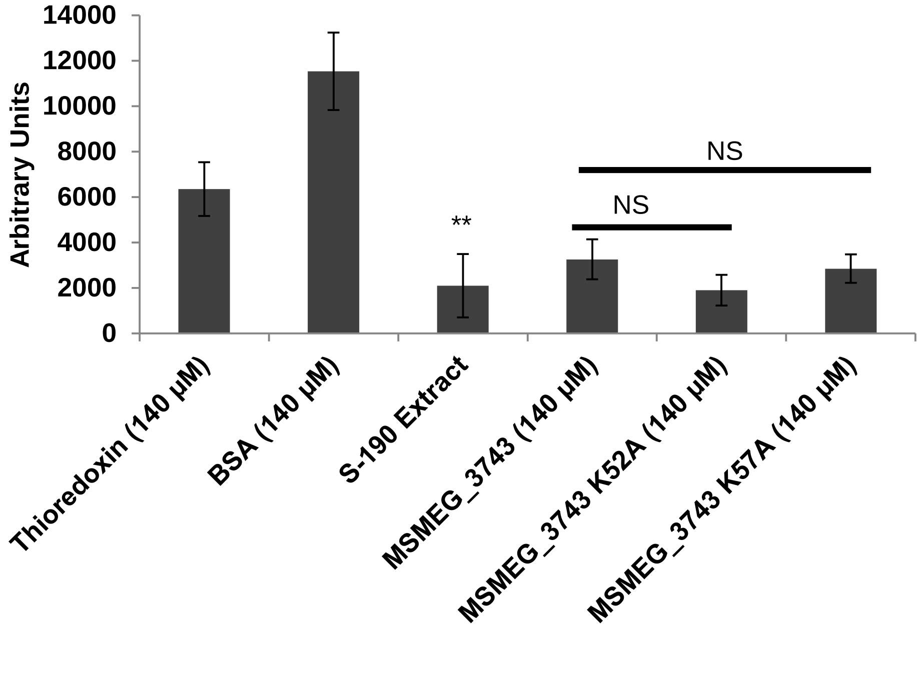


**Figure S3**: **ATPase activity of Lysine mutants of MSMEG_3743.** (a) Radiograph of γ-$^{32}$P-Pi release from hydrolysis of radiolabelled ATP. (b) Densitometric quantitation of unhydrolysed γ-$^{32}$P-ATP from each lane of the radiograph. Error bars represent SD; ** P<0.01, T-test done wrt. BSA. NS – Not Significant.

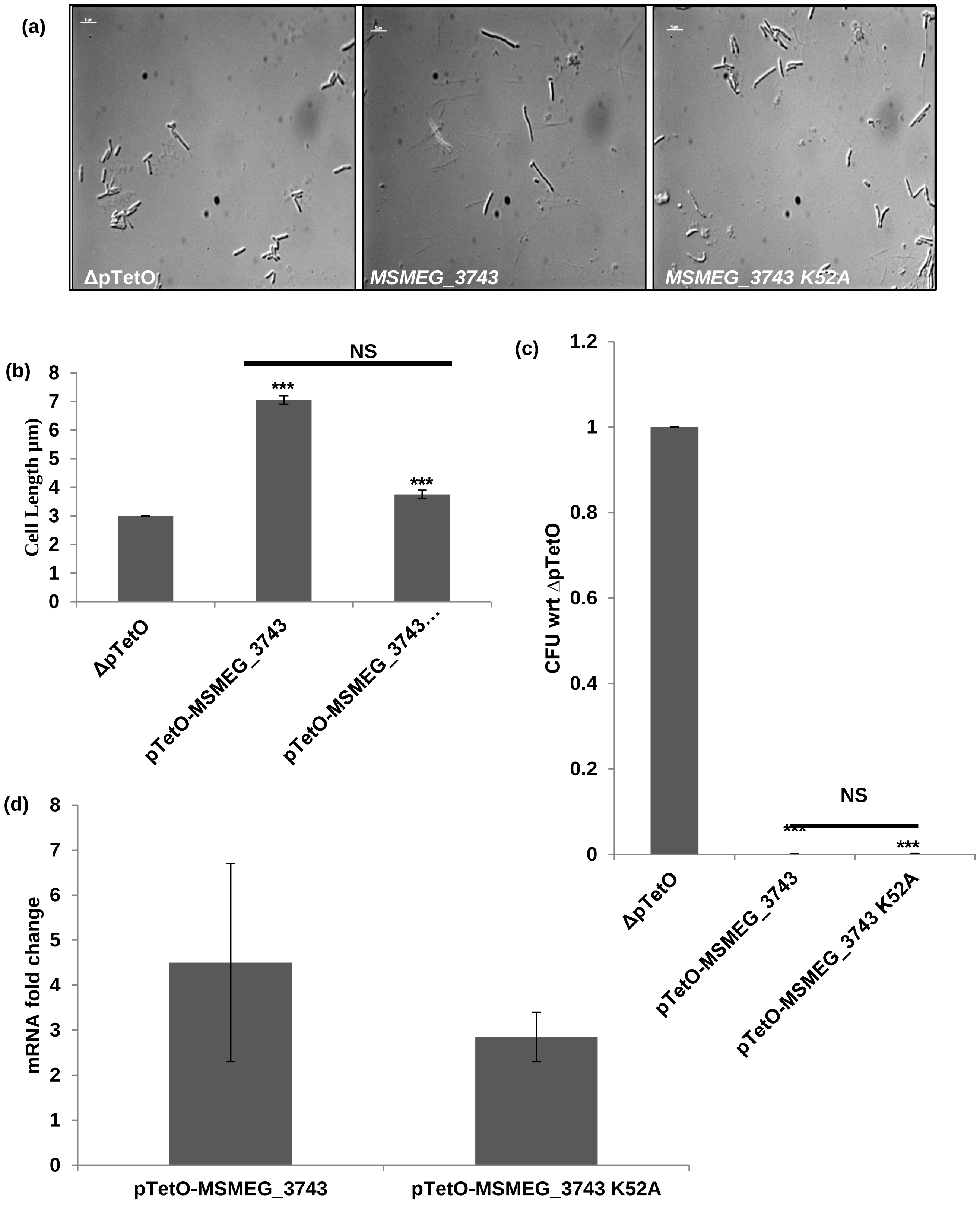


**Figure S4**: **Effect of overexpression of *MSMEG_3743 K52A* in *M. smegmatis*.** (a) Representative DIC images of overexpression strains; Scale bar 2 µm (b) Average cell lengths of overexpression strains (µm) Error bars represent SEM. (c) CFU counts, represented as a ratio wrt ΔpTetO (d) Transcript levels of *MSMEG_3743* & *MSMEG_3743* K52A following overexpression, wrt *Ms sigA*. *** P<0.001, T-test performed wrt ΔpTetO ; NS - Not Significant.

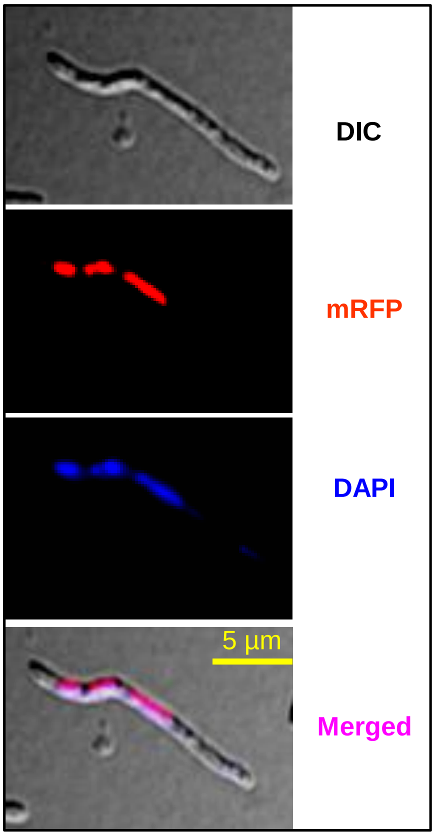


**Figure S5**: **Interaction of MSMEG_3743 with the bacterial chromosome.** Representative micrographs of *M. smegmatis* expressing *MSMEG_3743-mRFP* stained with DAPI; Scale bar 5 µm.

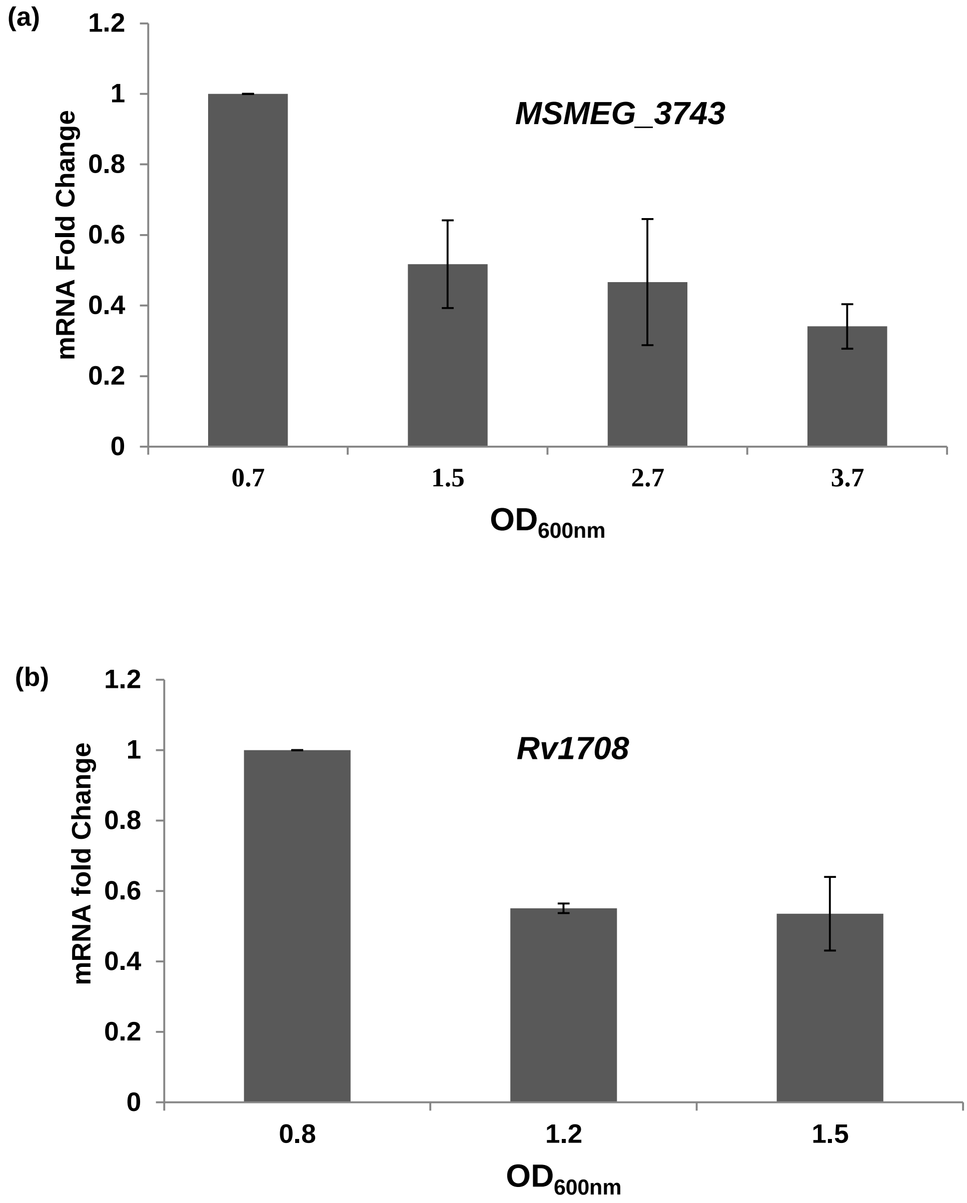


**Figure S6**: **Growth phase dependent gene expression of mycobacterial *minD* homologues.** Transcript levels of *MSMEG_3743* in *M. smegmatis* (a) and *Rv1708* in *M.tb* (b) at different phases of growth relative to their levels at mid-log phase. Error bars represent SD.

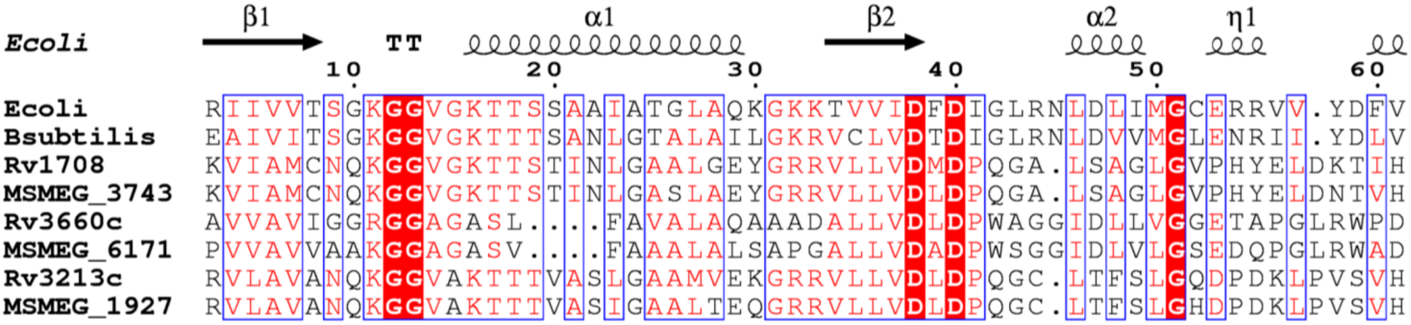

**Figure S7: Snippet of the sequence alignment containing the deviant Walker A motif (see figure S2) in *E.coli* MinD and its predicted homologues.** The two Glycine residues can be clearly seen to be conserved in all sequences.

| Primer Name | Description | Sequence (5'-3') |
|---|---|---|
| pTrc99A-*MSMEG_3743*-FP | FP (Forward Primer) for cloning *MSMEG_3743* in pTrc99A | ATGCAGAATTCATGCCTGACCAGCGGCGG |
| pTrc99A-*MSMEG_3743*-RP | RP (Reverse Primer) for cloning *MSMEG_3743* in pTrc99A | TGCATGGATCCTCACACGCCGAACCGGTG |
| pTrc99A- *MSMEG _6171*-FP | FP for cloning *Ms_6171* in pTrc99A | ATGCAGAATTCTTGCCCGTCATTGCCGGGC |
| pTrc99A- *MSMEG _6171*-RP | RP for cloning *Ms_6171* in pTrc99A | TGCATAAGCTTTCACGCCGCCTCCCGCAC |
| pTrc99A- *E. coli minD*-FP | FP for cloning *E. coli minD* in pTrc99A | ATGCAGAATTCATGGCACGCATTATTGTTG |
| pTrc99A- *E. coli minD*-RP | RP for cloning *E. coli minD* in pTrc99A | TGCATGGATCCTTATCCTCCGAACAAGCG |
| pTrc99A-*Rv1708*-FP | FP for cloning *Rv1708* in pTrc99A | ATGCAGAATTCTTGCCTGCGGGTCTCCCG |
| pTrc99A-*Rv1708*-RP | FP for cloning *Rv1708* in pTrc99A | TGCATAAGCTTTCACATGCCAAATCGGTCGA |
| pTrc99A-*Rv3660c*-FP | FP for cloning *Rv3660c* in pTrc99A | ATGCAGAATTCATGCTGACCGATCCGGGG |
| pTrc99A-*Rv3660c*-RP | RP for cloning *Rv3660c* in pTrc99A | TGCATGGATCCTCATGCCGCCCTACCGTG |
| pTetO-*MSMEG_3743*-FP | FP for cloning *MSMEG_3743* in pTetO | ATGCACATATGATGCCTGACCAGCGGCGG |
| pTetO-*MSMEG_3743*-RP | RP for cloning *MSMEG_3743* in pTetO | TGCATAAGCTTTCACACGCCGAACCGGTG |
| pTetO- *MSMEG _6171*-FP | FP for cloning *MSMEG _6171* in pTetO | ATGCACATATGTTGCCCGTCATTGCCGGGC |
| pTetO- *MSMEG _6*171-RP | RP for cloning *MSMEG _6171* in pTetO | TGCATAAGCTTTCACGCCGCCTCCCGCAC |
| pTetO- *Msm scpA*-FP | FP for cloning *Msm scpA* in pTetO | ATGCACATATGGTGAACGACGACGTGCGT |
| pTetO- *Msm scpA*-RP | RP for cloning *Msm scpA* in pTetO | TGCATAAGCTTCTATTCTTCCGCATCGGC |
| pTetO-*Msm scpB*-FP | FP for cloning *Msm scpB* in pTetO | ATGCACATATGATGACTGACGAGACCTCC |
| pTetO- *Msm scpB*-RP | RP for cloning *Msm scpB* in pTetO | TGCATAAGCTTTCAATCCTTGTCCACGTC |
| pTetO- *Msm parB*-FP | FP for cloning *Msm parB* in pTetO | ATGCACATATGATGAATCAGCCGGCACGC |
| pTetO- *Msm parB*-RP | RP for cloning *Msm parB* in pTetO | TGCATAAGCTTTTACTCGTTCTGGGCGCTC |
| pSCW54-*MSMEG_3743*-FP | FP for cloning *MSMEG_3743* in pSCW54 | ATGCACATATGATGCCTGACCAGCGGCGG |
| pSCW54-*MSMEG_3743*-RP | RP for cloning *MSMEG_3743* in pSCW54 | TGCATTTAATTAATCACACCGCCGAACCGGTG |
| pSCW54-6x His-*MSMEG_3743*-FP | FP for cloning N-terminal His-tagged *MSMEG_3743* in pSCW54 | ATGCACATATGATGCACCACCACCACCAC CACCCTGACCAGCGGCGGTCG |
| pSCW54-*MSMEG_3743*-6x His- RP | FP for cloning C-terminal His-tagged *MSMEG_3743* in pSCW54 | TGCATTTAATTAATCAGTGGTGGTGGTGGTG GTGCACGCCGAACCGGTG |
| pTetO-*MSMEG_3743*-NTL-FP | FP for cloning N-terminal mRFP fusion with *MSMEG_3743* in pSCW54 | ATGCAAAGCTTGGAGGAGGAGGAGGAATG CCTGACCAGCGGCGG |
| pTetO-*MSMEG_3743*-CTL-RP | RP for cloning N-terminal mRFP fusion with *MSMEG_3743* in pSCW54 | TGCATCATATGTCCTCCTCCTCCTCCCACGCCG AACCGGTGGAT |
| *MSMEG_3743*-RT-FP | FP for RT PCR of *MSMEG_3743* | TCGGCAAGACCACGTCGA |

| | | |
|---|---|---|
| *MSMEG_3743*-RT-RP | RP for RT PCR of *MSMEG_3743* | TCAGCACGTCGTCGATGG |
| *MSMEG _6171*-RT-FP | FP for RT PCR of *MSMEG _6171* | AGTCGTAAGGCCTGGCTG |
| *MSMEG _6171*-RT-RP | RP for RT PCR of *MSMEG _6171* | GCCACAAGATCGGTGTCC |
| *Msm sigA*-RT-FP | FP for RT PCR of *Msm sigA* | GCCAGCTCGGTGACTTCA |
| *Msm sigA*-RT-RP | RP for RT PCR of *Msm sigA* | CGTGACGCCGTAGACCTG |
| *Msm parB*-RT-FP | FP for RT PCR of *Msm parB* | CGAGTTCGGTCTCATGCAG |
| *Msm parB*-RT-RP | RP for RT PCR of *Msm parB* | GTTCAACTGGACGCGGTG |
| *Rv1708*-RT-FP | FP for RT PCR of *Rv1708* | ATGGATCCGCAAGGAGCG |
| *Rv1708*-RT-RP | RP for RT PCR of *Rv1708* | ACCCACCTCGTTGACCAG |
| *Mtb. sigA*-RT-FP | FP for RT PCR of *Mtb. sigA* | AAACAGATCGGCAAGGTAGC |
| *Mtb. sigA*-RT-RP | RP for RT PCR of *Mtb. sigA* | TCCAGCGATGGTTTTTCG |
| pET22b-*MSMEG_3743*-FP | FP for cloning *MSMEG_3743* in pET22b | GGAATTCCATATGCCTGACCAGCGGCGGTC |
| pET22b-*MSMEG_3743*-RP | RP for cloning *MSMEG_3743* in pET22b | CGG CTCGAG CACGCCGAACCGGTGGATG |
| *MSMEG_3743* K52A-FP | FP for SDM PCR of *MSMEG_3743K52A* | ATGTGCAACCAGGCGGGCGGC |
| *MSMEG_3743* K52A-RP | RP for SDM PCR of *MSMEG_3743K52A* | GCCGACGCCGCCCGCCTGGTTG |
| *MSMEG_3743* K57A-FP | FP for SDM PCR of *MSMEG_3743K57A* | GCGGCGTCGGCGCGACCACGT |
| *MSMEG_3743* K57A-RP | RP for SDM PCR of *MSMEG_3743K57A* | GTCGACGTGGTCGCGCCGACG |
| pJEX55-*MSMEG_3743*-FP | FP for cloning *MSMEG_3743* in pJEX55 | ATGCAGGATCCATGCCTGACCAGCGGCGG |
| pJEX55-*MSMEG_3743*-RP | RP for cloning *MSMEG_3743* in pJEX55 | TGCATGAATTCTCACACGCCGAACCGGTG |
| pUAB400-*MSMEG_3743*-FP | FP for cloning *MSMEG_3743* in pUAB400 | CCGGAATTCGTGGGCCTGACGGGCCGG |
| pUAB400-*MSMEG_3743*-RP | RP for cloning *MSMEG_3743* in pUAB400 | CCCAAGCTTTCACACGCCGAACCGGTGGA |
| pUAB300- *MSMEG parB*-FP | FP for cloning *MSMEG parB* in pUAB300 | CGCGGATCCATGAATCAGCCGGCACGC |
| pUAB300- *MSMEG parB*-RP | RP for cloning *MSMEG parB* in pUAB300 | CCCAAGCTTTTACTCGTTCTGGGCGCT |
| pUAB400-*Rv1708*-FP | FP for cloning *Rv1708* in pUAB400 | GGAATTCTTGCCTGCGGGTCTCCCG |
| pUAB400-*Rv1708*-RP | RP for cloning *Rv1708* in pUAB400 | CCCAAGCTTTCACATGCCAAATCGGTCGATC |
| pUAB300-*Mtb. parB*-FP | FP for cloning *Mtb. parB* in pUAB300 | GGAAGATCTCATGACCCAGCCGTCACG |

| | | |
|---|---|---|
| pUAB300-*Mtb. parB*-RP | RP for cloning *Mtb. parB* in pUAB300 | CCCAAGCTTTTACAGAGCGTCCCTGTGC |
| pUAB300-*Mtb. scpA*-FP | FP for cloning *Mtb. scpA* in pUAB300 | CGGGATCCGTGAACGGCCTTCAGAAC |
| pUAB300-*Mtb. scpA*-RP | RP for cloning *Mtb. scpA* in pUAB300 | CCCAAGCTTTCACAAGCGCCGCTCCTT |
| pUAB300-*Mtb. scpB*-FP | FP for cloning *Mtb. scpB* in pUAB300 | CGGGATCCGTGACCGAACATATGCCC |
| pUAB300-*Mtb. scpB*-RP | RP for cloning *Mtb. scpB* in pUAB300 | CCCAAGCTTTCAATCACGGTCCACGTC |
| pUAB300-*Rv1707*-FP | FP for cloning *Rv1707* in pUAB300 | GGAAGATCTGTGTTACAACGAATCGCTAG |
| pUAB300-*Rv1707*-RP | RP for cloning *Rv1707* in pUAB300 | CCCAAGCTTTCAGGCGGATTCGAGGAC |
| pGEX6P1-*Rv1708*-FP | FP for cloning *Rv1708* in pGEX6P1 | GGAATTCTTGCCTGCGGGTCTCCCG |
| pGEX6P1-*Rv1708*-RP | RP for cloning *Rv1708* in pGEX6P1 | CCGCTCGAGTCACATGCCAAATCGGTCGATC |
| pET22b-*Mtb. scpA*-FP | FP for cloning *Mtb. scpA* in pET22b | GGAATTCCATATGGTGAACGGCCTTCAGAACGA |
| pET22b-*Mtb. scpA*-RP | RP for cloning *Mtb. scpA* in pET22b | CCGCTCAGCAAGCGCCGCTCCTTCTC |
| pET22b-*Mtb. parB*-FP | FP for cloning *Mtb. parB* in pET22b | GGAATTCCATATGACCCAGCCGTCACGCAGA |
| pET22b-*Mtb. parB*-RP | RP for cloning *Mtb. parB* in pET22b | CCCAAGCTTCAGAGCGTCCCTGTGAAG |

**Table S1: Oligonucleotides used in this study**
